# ALKBH7 mediates necrosis via rewiring of glyoxal metabolism

**DOI:** 10.1101/2020.05.04.077297

**Authors:** Chaitanya A. Kulkarni, Sergiy M. Nadtochiy, Leslie Kennedy, Jimmy Zhang, Sophea Chhim, Hanan Alwaseem, Elizabeth Murphy, Dragony Fu, Paul S. Brookes

## Abstract

Alkb homolog 7 (ALKBH7) is a mitochondrial α-ketoglutarate dioxygenase required for necrotic cell death in response to DNA alkylating agents, but its physiologic role within tissues remains unclear. Herein, we show that ALKBH7 plays a key role in the regulation of dialdehyde metabolism, which impacts cardiac survival in response to ischemia-reperfusion (IR) injury. Using a multi-omics approach, we do not find evidence that ALKBH7 functions as a prolyl-hydroxylase. However, we do find that mice lacking ALKBH7 exhibit a significant increase in glyoxalase I (GLO-1), a dialdehyde detoxifying enzyme. Consistent with increased dialdehyde production, metabolomics analysis reveals rewiring of metabolic pathways related to the toxic glycolytic by-product methylglyoxal (MGO), as well as accelerated glycolysis and elevated levels of MGO protein adducts, in mice lacking ALKBH7. Consistent with roles for both necrosis and glycative stress in cardiac IR injury, hearts from male but not female *Alkbh7*^-/-^ mice are protected against IR, although somewhat unexpectedly this protection does not appear to involve modulation of the mitochondrial permeability transition pore. Highlighting the importance of MGO metabolism for the observed protection, removal of glucose as a metabolic substrate or pharmacologic inhibition of GLO-1 both abrogate cardioprotection in ALKBH7 deficient mice. Integrating these observations, we propose that ALKBH7 plays a role in the regulation of glyoxal metabolism, and that protection against necrosis and IR injury bought on by ALKBH7 deficiency originates from hormetic signaling in response to elevated MGO stress.

## INTRODUCTION

The α-ketoglutarate (α-KG) dioxygenases are a diverse enzyme superfamily, whose primary biochemical function is the addition of hydroxyl (–OH) to protein or nucleic acid substrates.^1^ The family includes the TET 5-methylcytosine hydroxylases, the EGLN prolyl-hydroxylases that regulate hypoxia inducible factor (HIF), and the JmjC domain-containing histone demethylases. All α-KG dioxygenases use α-KG and O_2_ as biochemical substrates and generate succinate as product. The AlkB homologs (ALKBHs) are a distinct family of nine α-KG dioxygenases that are homologs of *E. coli* AlkB.^2^ The bacterial AlkB enzyme catalyzes demethylation of DNA damaged by alkylating agents, via hydroxylation of the methylated DNA followed by spontaneous decomposition to release formaldehyde and recover the DNA base.^3,4^ Many eukaryotic ALKBHs have been shown to act on DNA or RNA substrates, including mammalian ALKBHs 1-3, 5, 8 and FTO.^5-8^ The subject of this investigation is ALKBH7, a poorly characterized mitochondrial α-KG dioxygenase which has no known substrates.

In contrast to a role in DNA repair or RNA modification, structural studies have revealed ALKBH7 lacks a critical nucleotide recognition lid required for binding DNA or RNA.^9^ Moreover, A study of mitochondria from several tissues of *Alkbh7*^-/-^ mice showed no differences in mtDNA modifications (6-methyladenine, 5-methylcytosine, etc.) vs. wild-type (WT) at young ages, although *Alkbh7*^-/-^ mtDNA did accumulate more modifications in old-age.^10^ Together these observations suggest ALKBH7 may not play significant roles in either the repair of known AlkB substrates, or in oxidizing as-yet unknown nucleic acid substrates. Similarly, efforts to identify potential protein substrates of ALKBH7 have not yielded insight to its function, with both a yeast-2-hybrid screen and several large mitochondrial protein:protein interaction databases not reporting any ALKBH7-binding proteins.^11-14^

While the nucleic acid or protein substrate(s) of ALKBH7 remain(s) unclear, previous studies using RNAi or genetic ablation in human cells have found that ALKBH7 is required for programmed necrosis induced by DNA alkylating agents.^15^ Moreover, *Alkbh7*^-/-^ mice exhibit protection against alkylation-induced cell death in certain tissues.^16^ Notably this phenotype is only observed in males, even though *Alkbh7* is not a sex-linked gene. Furthermore, the *Alkbh7*^-/-^ mouse exhibits obesity due to defective fatty acid β-oxidation,^17^ and an *Alkbh7* mis-sense mutation (R191Q) has been linked to prostate cancer.^18^ However, it is unknown how these phenotypes are linked to the biochemical function of ALKBH7 as an α-KG dioxygenase.

The heart is a mitochondria-rich tissue, and cardiomyocyte necrosis plays a key role in cardiac pathology such as that occurring in ischemia-reperfusion (IR) injury.^19^ As such, the requirement for ALKBH7 in other models of necrosis^15^ makes the protein a potential target for the modulation of cell death in response to IR injury. Herein, focusing on heart tissue we employed a multi-omics approach to elucidate ALKBH7 biology, finding that hearts from *Alkbh7*^-/-^ mice are protected against IR injury. We also find that a core component of this protected phenotype is hormetic signaling, resulting in a rewiring of metabolism in response to elevated glycative stress. These findings imply potential therapeutic utility for ALKBH7 inhibitors to prevent necrosis in IR and other conditions.

## RESULTS

The complete original data used to generate all figures in the main document and supplement are contained in a spreadsheet available at DOI: 10.6084/m9.figshare.12200273 (DOI reserved, unembargoed upon publication).

### Proteomic analysis to identify ALKBH7 substrates suggests it is not a prolyl-hydroxylase

Several members of the α-KG dioxygenase superfamily possess prolyl-hydroxylase activity, and ALKBH7 is known to auto-hydroxylate on Leucine 110, suggesting it has hydroxylase activity.^9^ To investigate the hypothesis that ALKBH7 may be a prolyl-hydroxylase, a tandem-mass-tag (TMT) proteomic approach was employed to identify potential targets, assuming such targets would contain less hydroxyproline (P-OH) in *Alkbh7*^-/-^ vs. WT samples. Using heart tissue from *Ablkbh7*^-/-^ and WT mice (**Fig. S1**), a total of 451 P-OH containing peptides were identified, and their abundances normalized to those of their 238 parent proteins. Differential analysis, applying thresholds of ±1.5-fold change and p<0.05 for significance, revealed only a handful of peptides with altered P-OH (volcano plot **Fig. 1A**, top 5 up/down hits in **Fig. 1B**). Only one peptide showed significantly less P-OH: the β-oxidation enzyme hydroxyacyl-CoA dehydrogenase (*Hadh* gene).

**Figure 1.**
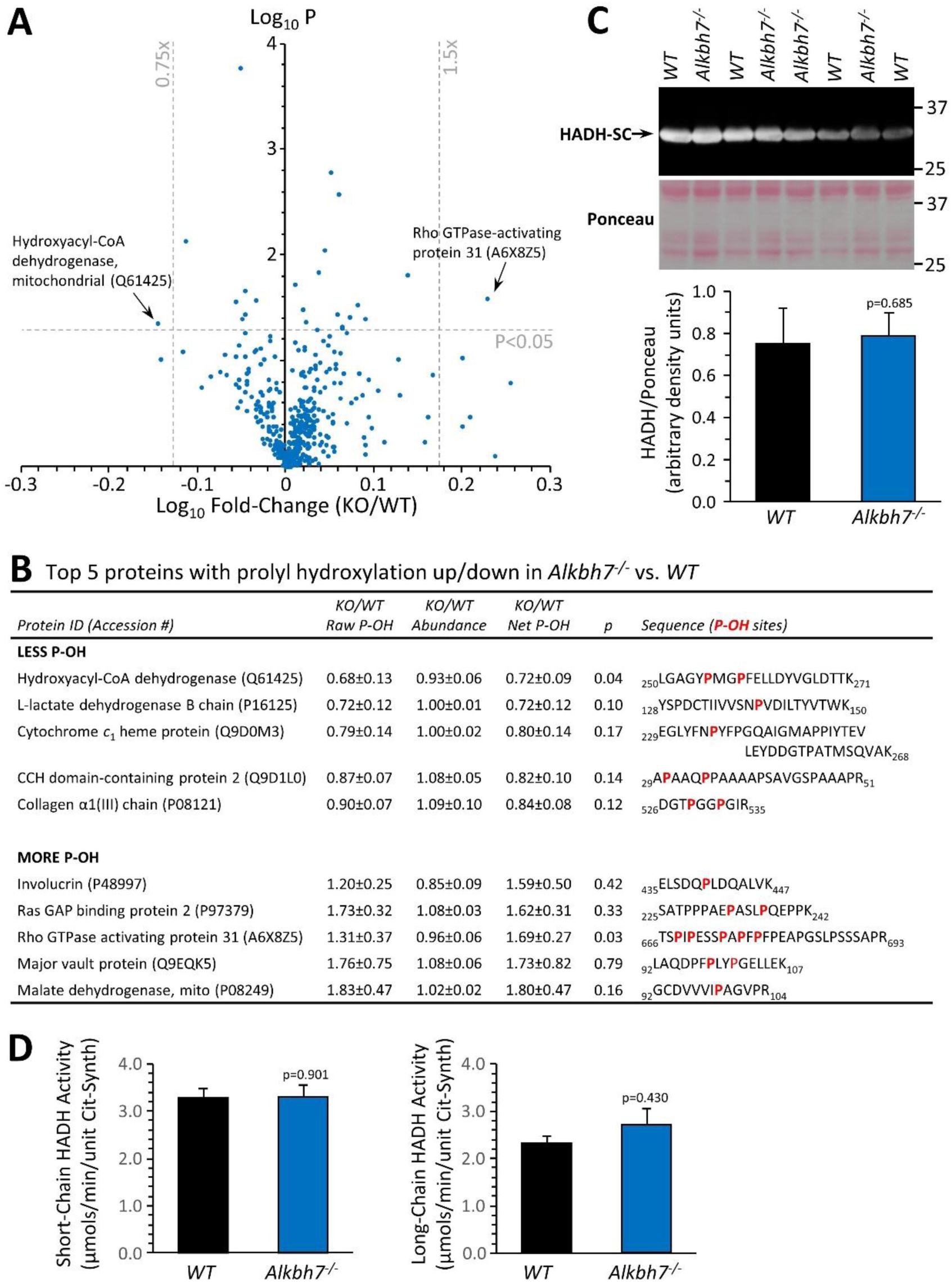
Proteomic analysis of prolyl-hydroxylation in WT vs. *Alkbh7*^-/-^. Hearts from young male WT or *Alkbh7*^-/-^ mice were analyzed by tandem mass tag LC-MS/MS as per the methods. Abundance of each P-OH peptide was normalized to the abundance of its parent protein. **(A):** Volcano plot showing relative levels of 451 P-OH containing peptides. X-axis shows Log_10_ of fold change (*Alkbh7*^-/-^ / WT) and Y-axis shows Log_10_ of significance (paired t-test, N=5). Proteins crossing thresholds (gray lines) in upper left or right quadrants are labeled. **(B):** Table showing the top 5 P-OH containing peptides exhibiting increased or decreased relative abundance in *Alkbh7*^-/-^ vs. WT. Table shows raw abundance of each P-OH peptide, abundance of the parent protein, and normalized abundance of the P-OH peptide. Annotated sequences highlight the hydroxylated proline residues in red. **(C):** Western blot showing abundance of HADH-SC (*Hadh*) in WT or *Alkbh7*^-/-^ heart mitochondria with quantitation below, normalized to protein loading determined by Ponceau S stained membrane. **(D):** Spectrophotometric activity assays of short-chain and long-chain HADH in WT or *Alkbh7*^-/-^ heart mitochondria. Bar graphs in panels C/D show means ± SE, N=3-5, with p-values (paired t-test) shown above error bars.

It is reported that *Alkbh7*^-/-^ mice are obese and harbor a baseline defect in β-oxidation of long-chain fatty acids such as oleate, which can be overcome when stimulated by fasting.^17^ To determine β-oxidation levels in the heart, a mostly fat-burning organ, Seahorse XF analysis of isolated cardiomyocytes from *Alkbh7*^-/-^ and WT mice was undertaken, revealing a small but significant decrease in oleate oxidation at baseline, with this effect disappearing upon stimulation of maximal respiration (**Fig. S2**). Although HADH is primarily involved in the β-oxidation of short chain fatty acids, we hypothesized based on P-OH proteomic data and the obese phenotype that prolyl-hydroxylation of HADH may regulate its activity. However, western blotting showed no alteration in HADH protein levels between WT and *Alkbh7*^-/-^ (**Fig. 1B**), and activity assays of both short-chain HADH (*Hadh* gene) and long-chain HADH (*Hadha* gene) revealed no differences between genotypes (**Fig. 1D**). As such, we consider it unlikely that prolyl-hydroxylation by ALKBH7 is an underlying cause of defective β-oxidation in the *Alkbh7*^-/-^ mouse.

To identify potential ALKBH7 binding partners, a FLAG-tag pull-down interactome experiment was performed, under either baseline or DNA alkylation stress conditions. As **Table S1** shows, several mitochondrial heat shock proteins were identified as ALKBH7 interactors, despite no such proteins being differentially hydroxylated (**Fig. 1A**). This finding is in agreement with a recent antibody-based immunoprecipitation study which suggested a role for ALKBH7 in proteostasis,^20^ although the functional significance of this for necrosis is unclear (see **Fig. S8** and related text). An additional protein hit was the NDUFS7 subunit of respiratory complex I, which the BioPlex interactome database also reports as an ALKBH7 interacting protein.^21^ However, enzyme assays in heart and liver mitochondria from WT and *Alkbh7*^-/-^ mice revealed no differences in the activities of complex I and several other key mitochondrial enzymes (**Fig. S3**), suggesting no role for ALKBH7 in regulating complex I function. Overall, consistent with a general paucity of ALKBH7 binding proteins,^11-14^ we consider it unlikely that the necrosis function of ALKBH7 is due to prolyl-hydroxylation or the binding and modulation of mitochondrial heat shock proteins or respiratory complexes.

### Proteomic abundance analysis in Alkbh7^-/-^ indicates re-wiring of glyoxal metabolism

In parallel with analysis of P-OH, the TMT proteomic experiment also yielded relative abundance values for 3,737 proteins in *Alkbh7*^-/-^ and WT hearts, with a volcano plot (**Fig. 2A**) revealing several differences which may underlie the metabolic phenotype of the knockout animals.^A^ The lipid droplet protein perilipin-5, which signals via Sirt1/PPAR-α to drive mitochondrial biogenesis and fat oxidation,^22^ was 25% lower in *Alkbh7*^-/-^ vs. WT. In addition, fructose-1,6-bisphosphatase 2 (FBP2/PFK2) was 31% lower in *Alkbh7*^-/-^ vs. WT, a finding typically associated with acceleration of glycolysis.^23^

**Figure 2.**
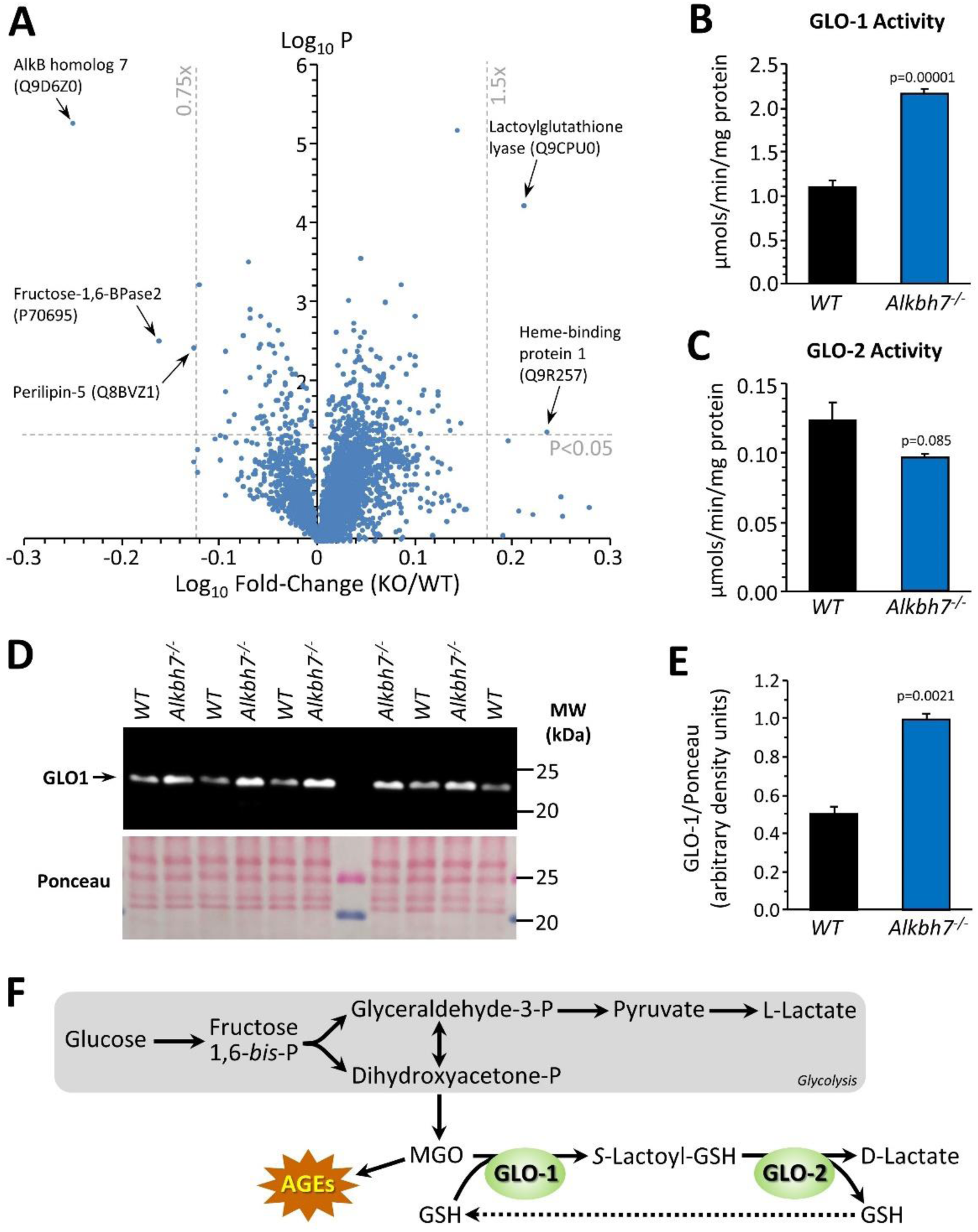
Proteomic analysis of protein abundance in WT vs. *Alkbh7*^-/-^. Hearts from young male WT or *Alkbh7*^-/-^ mice were analyzed by tandem mass tag LC-MS/MS as per the methods. **(A):** Volcano plot showing relative levels of 3,737 proteins. X-axis shows Log_10_ of fold change (*Alkbh7*^-/-^ / WT) and Y-axis shows Log_10_ of significance (paired t-test, N=5). Proteins crossing thresholds (gray lines) in upper left or right quadrants are labeled. **(B):** Activity of GLO-1 in WT or *Alkbh7*^-/-^ heart cytosol. **(C):** Activity of GLO-2 in WT or *Alkbh7*^-/-^ heart cytosol. **(D):** Western blot showing abundance of GLO-1 in WT or *Alkbh7*^-/-^ heart cytosol, with Ponceau stained membrane below. **(E)**: Quantitation of GLO-1 blot, normalized to protein loading. Bar graphs in panels B/C/E show means ± SE, N=4-5, with p-values (paired t-test) shown above error bars. **(F):** Schematic showing the methylglyoxal detoxification system and its relationship to glycolysis. Abbreviations: AGEs: Advanced glycation end products, GSH: glutathione. MGO: methylglyoxal.

In addition, a highly significant (p=0.00006) 1.6-fold elevation was seen in lactoylglutathione lyase (glyoxalase I, GLO-1) in *Alkbh7*^-/-^ vs. WT. This observation was confirmed by enzymatic activity assay (**Fig. 2B**) and by western blot (**Fig. 2D/E**), with a similar activity difference also observed in *Alkbh7*^-/-^ vs. WT liver tissue (**Fig. S3**). GLO-1 is part of the dialdehyde detoxification pathway that handles toxic metabolites such as the glycolytic by-product methylglyoxal (MGO), recycling it to D-lactate, thus avoiding the generation of advanced glycation end products (**Fig. 2F**).^24^ No change was seen in the activity of the companion enzyme GLO-2 in *Alkbh7*^-/-^ (**Fig. 2C**). The only other protein significantly upregulated in *Alkbh7*^-/-^ was heme binding protein 1 (Hebp1), and notably a recent study found both GLO-1 and Hebp1 were upregulated in Alzheimer’s disease,^25^ suggesting these proteins may share a common upstream regulator. Overall, despite extensive proteome coverage, a surprisingly small number of proteins (4) were up- or down-regulated in *Alkbh7*^-/-^ heart.

Although ALKBH7 is generally thought to be mitochondrial, several of the differences observed between WT and *Alkbh7*^-/-^ heart were cytosolic proteins, including GLO-1. In this regard, western blotting (**Fig. S1**) showed immunoreactivity for ALKBH7 in the cytosolic compartment (uncontaminated by the mitochondrial marker ANT-1) as well as in mitochondria, suggesting ALKBH7 may not be exclusively mitochondrial. The relative importance of these sub-populations of ALKBH7 in driving necrosis or other phenotypes is currently unclear.

### Metabolomics analysis in Alkbh7^-/-^ confirms rewired glyoxal metabolism

To further probe metabolism in *Alkbh7*^-/-^ hearts, a steady-state metabolomics analysis was undertaken (**Fig. 3A**), which revealed perturbations in several key metabolites related to MGO stress. The antioxidants carnosine and glutathione (GSH) were both significantly lower in *Alkbh7*^-/-^, consistent with their being utilized in the detoxification of MGO.^26^ As noted above (**Fig. 2F**), GLO-2 recycles GSH consumed by GLO-1, so an elevation in GLO-1 activity without a concomitant upregulation of GLO-2 would be predicted to result in GSH depletion. Furthermore, numerous metabolites in the lower half of glycolysis were elevated in *Alkbh7*^-/-^, suggesting acceleration of this pathway (**Fig. 3B**). To test this hypothesis directly, ^13^C-glucose tracing was employed to measure glycolytic flux in perfused mouse hearts,^27^ and the results in **Fig. 3C** show that glycolytic flux was indeed faster in *Alkbh7*^-/-^. Further evidence for elevated MGO stress in *Alkbh7*^-/-^ was seen in the form of elevated MGO protein adduct levels in both cytosol and mitochondria (**Fig. 3D-F**). Together these data suggest the *Alkbh7*^-/-^ heart experiences elevated glycative stress, potentially driving a hormetic response.

**Figure 3.**
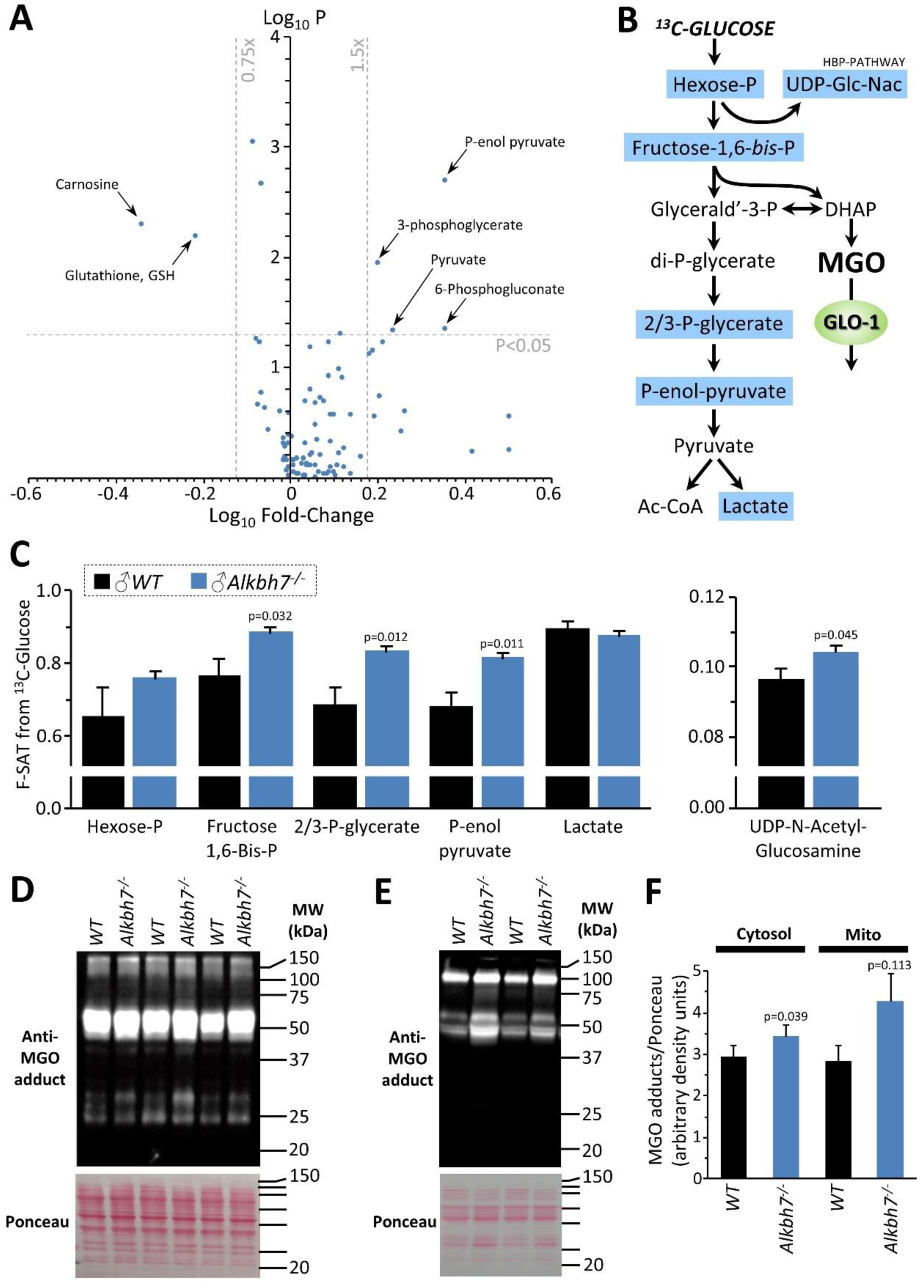
Metabolomics analysis in WT vs. *Alkbh7*^-/-^. Hearts from young male WT or *Alkbh7*^-/-^ mice were analyzed by LC-MS/MS based metabolomics as per the methods. **(A):** Volcano plot showing relative levels of 90 metabolites in the steady state. X-axis shows Log_10_ of fold change (*Alkbh7*^-/-^ / WT) and Y-axis shows Log_10_ of significance (paired t-test, N=8-17 depending on metabolite). Metabolites crossing thresholds (gray lines) in upper left or right quadrants are labeled. A pathway impact analysis is shown in Fig. S4. **(B):** Schematic showing glycolysis and its relationship to methylglyoxal (MGO). Metabolites quantified in ^13^C-flux measurements (panel C) are highlighted blue. **(C):** ^13^C-glucose flux measurements of glycolytic activity in *Alkbh7*^-/-^ vs. WT hearts. Y-axis shows fractional saturation (F-SAT) of each metabolite within 5 min. from exogenously delivered [U-^13^C] glucose. Note: UDP-Glc-Nac is shown on separate axes for clarity. **(D/E):** Western blot showing abundance of MGO-adducts in *Alkbh7*^-/-^ and WT heart cytosol (D) or mitochondria (E). Ponceau stained membranes are shown below. **(F):** Quantitation of MGO adduct content from blots, normalized to protein loading. Bar graphs in panels C/F show means ± SE, N=4-5, with p-values (paired t-test) shown above error bars.

### Loss of ALKBH7 protects the heart from ischemia-reperfusion (IR) injury

In addition to metabolic effects, a key phenotype resulting from *Alkbh7* ablation is protection against necrosis.^15^ In seeking links between glyoxal metabolism and necrosis, it is notable that both glycative stress and necrosis are implicated in the pathology of cardiac IR injury.^28-30^ In addition, a mitochondrially-targeted MGO scavenging molecule was recently shown to protect the heart against IR injury.^31^ To test the hypothesis that loss of ALKBH7 may protect against IR, perfused hearts from WT and *Alkbh7*^-/-^ mice were subjected to IR injury (25 min. global ischemia, 60 min. reperfusion). As shown in **Fig. 4A/B**, male *Alkbh7*^-/-^ hearts exhibited significantly improved post-ischemic functional recovery and significantly lower infarct size (an indicator of necrosis) compared to WT. Consistent with sexual dimorphism in the necrosis effects of ALKBH7,^16^ no protection against IR injury was observed in hearts from female *Alkbh7*^*-*/-^ mice (**Fig. S5**).

**Figure 4.**
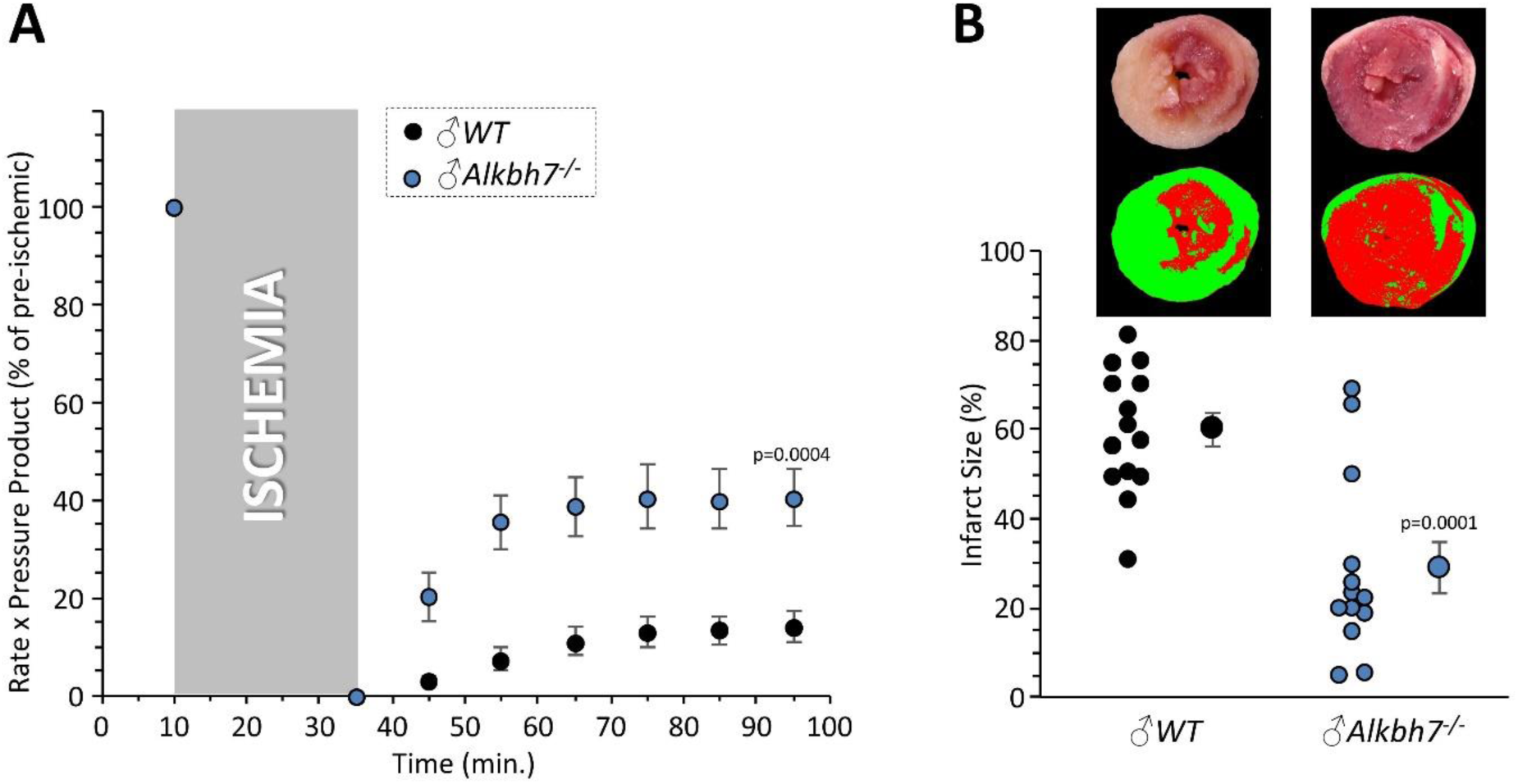
Response to ex-vivo cardiac ischemia-reperfusion (IR) injury in WT vs. *Alkbh7*^-/-^. Hearts from young male WT and *Alkbh7*^-/-^ mice were Langendorff perfused and subjected to 25 min. ischemia plus 60 min. reperfusion. **(A):** Cardiac function assessed by left ventricular balloon pressure transducer. Graph shows the product of heart rate multiplied by left ventricular developed pressure, as a percentage of the initial (pre-ischemic) value. **(B):** Post IR staining with TTC for quantitation of myocardial infarct size. Representative TTC-stained heart slices are shown, with pseudo-colored mask images used for quantitation by planimetry (red = live tissue, green = infarct). Data are quantified below, with individual data points to shown N, and means ± SE. p-values (paired t-test) are shown above error bars.

Both hormetic signaling (e.g., via *Nrf-2*) and several paradigms of cardioprotection are known to decline with age. ^32-35^ In addition GLO-1 activity declines with age,^24^ and the *Glo1* gene contains an Nrf-2 inducible antioxidant response element.^36^ As such, agreeing with the observed cardioprotection in the *Alkbh7*^-/-^ mouse originating via hormetic signaling, we also found that protection against IR injury was lost in aged male *Alkbh7*^-/-^ mice (**Fig. S6**). Together, these results suggest that ALKBH7 may play a role in necrosis by regulating MGO metabolism, and its ablation triggers a hormetic signaling response that endows protection against IR in the heart.

### Acute pharmacologic ALKBH inhibition elicits cardioprotection

The activity of α-KG dioxygenases can be inhibited by D- or L-isomers of the non-canonical metabolite 2-hydroxyglutarate (2-HG),^37^ with L-2-HG inhibiting ALKBHs more potently than D-2-HG.^38,39^ In addition, acute administration of the generic α-KG dioxygenase inhibitor dimethyloxalylglycine was shown to confer protection against hypoxic injury in cardiomyocyte model of IR.^40^ Since genetic ablation of ALKBH7 was cardioprotective, we thus hypothesized its pharmacologic inhibition may serve a similar purpose. As **Fig. S7** shows, administration of L-2-HG as its dimethyl ester (a common delivery strategy for dicarboxylates) was cardioprotective in WT hearts, eliciting enhanced functional recovery and lower infarct size (albeit the latter non-significant). While L-2-HG is known to have multiple targets, these data suggest the development of more specific ALKBH7 inhibitors may be a promising therapeutic strategy for IR injury.

### Cardioprotection in Alkbh7^-/-^ is not due to the mitochondrial unfolded protein response

We recently showed that activation of the mitochondrial unfolded protein response (UPR^mt^) is sufficient to induce cardioprotection against IR injury.^41^ The genes encoding *LonP1* and *ClpP*, two mitochondrial proteases involved in UPR^mt^ signaling,^42^ are also located on mouse chromosome 17 adjacent to the *Alkbh7* gene. In addition, a recent proteomic study proposed a role for ALKBH7 in mitochondrial proteostasis,^20^ and our pull-down experiment identified several mitochondrial heat-shock proteins as potential ALKBH7 interactors (**Table S1**). Furthermore, the related protein ALKBH1 has been shown to partially localize to mitochondria, and its knock-down induces a UPR^mt^.^43^ As such, we hypothesized constitutive UPR^mt^ activation might underlie the cardioprotective effects of ALKBH7 ablation. However, western blotting (**Fig. S8**) showed only small increases in LonP1 and ClpP protein in *Alkbh7*^-/-^ (the former non-significant), and a significant decrease in HSP60 (*Hspd1*) protein. We also did not find any UPR^mt^ target proteins upregulated in our proteomics analysis (**Fig. 2**). Overall these observations suggest that modulation of the UPR^mt^ is not a key mechanism by which ALKBH7 regulates necrosis.

### Cardioprotection in Alkbh7^-/-^ is not via the mitochondrial permeability transition pore

A core component of the necrotic cell death machinery is the mitochondrial permeability transition (PT) pore, which is regulated by the cis/trans prolyl-isomerase cyclophilin D (CypD, *ppif*).^44^ Parallels between CypD and ALKBH7 function have previously been speculated.^9^ In addition, although somewhat counter-intuitive, it has been shown that MGO can inhibit the PT pore,^45^ and our data suggest *Alkbh7*^-/-^ mice experience greater MGO stress (**Figs. 2,3**). Thus, we hypothesized ALKBH7 may regulate the PT pore. However, an osmotic swelling PT pore assay in isolated cardiac mitochondria from WT and *Alkbh7*^-/-^ mice revealed only a slight blunting of pore opening in *Alkbh7*^-/-^ (**Fig. 5A/B**). In addition, pore opening in both genotypes was inhibited by CypD inhibitor cyclosporin A, suggesting no differences in the underlying ability of CypD to regulate the pore. An isolated mitochondrial Ca^2+^ handling assay (**Fig. 5C-E**) showed a slight elevation in the amount of Ca^2+^ required to trigger the pore in *Alkbh7*^-/-^, and no difference in Ca^2+^ uptake kinetics. Furthermore, blue-native gel analysis of ATP synthase multimers, which are postulated to contribute to the composition of the PT pore,^46^ showed no differences between *Alkbh7*^-/-^ and WT (**Fig. S9**). Together these findings suggest the mitochondrial PT pore is not a central mechanism by which ALKBH7 regulates necrosis.

**Figure 5.**
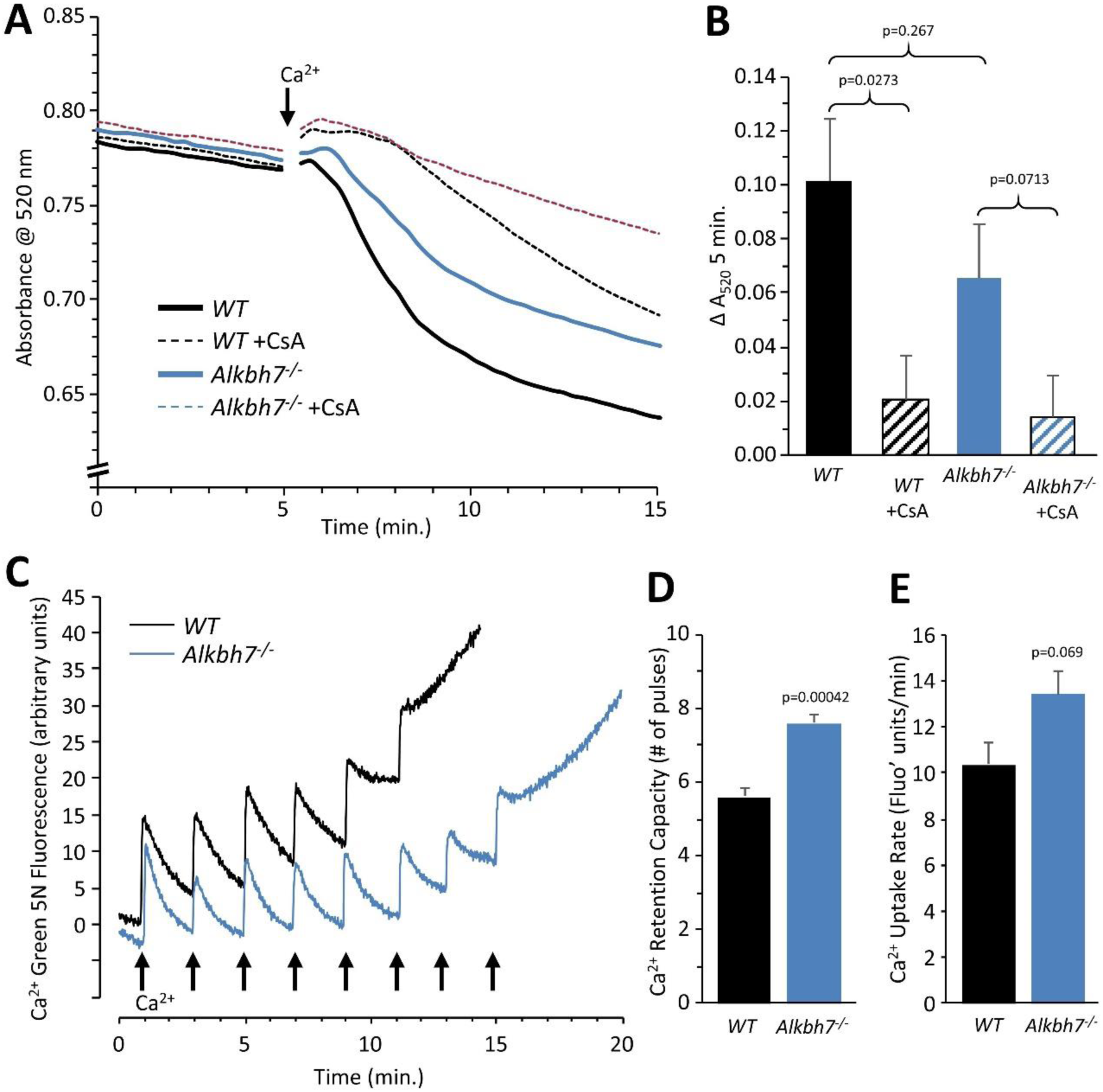
Mitochondrial PT pore & Ca^2+^ handling in WT vs. *Alkbh7*^-/-^. **(A):** Opening of the mitochondrial PT pore was assayed spectrophotometrically in isolated cardiac mitochondria from young male WT and *Alkbh7*^-/-^ mice. Average traces are shown, with addition of 100 µM Ca^2+^ to initiate PT pore opening and swelling indicated by the arrow. Dotted lines indicate the presence of the PT pore inhibitor cyclosporin A (CsA). Error bars are omitted for clarity. **(B):** Quantitation of pore opening, as the change in swelling (absorbance at 520 nm) in 5 min. Data are means ± SE, N=7, with significance between groups (unpaired t-test) shown above error bars. **(C):** Mitochondrial Ca^2+^ handling assayed by Ca^2+^ green-5N fluorescence. Isolated cardiac mitochondria from young male WT and *Alkbh7*^-/-^ mice were incubated with Ca^2+^ green-5N to indicate extra-mitochondrial [Ca^2+^]. Pulses of 10 µM Ca^2+^ were added at ∼2 min. intervals as indicated by arrows. Representative traces are shown. **(D):** Quantitation of the number of Ca^2+^ pulses tolerated by mitochondria before PT pore opening occurred (as indicated by a sharp upward deflection in the Ca^2+^ green-5N trace). **(E):** Quantitation of the initial rate of mitochondrial Ca^2+^ uptake, calculated from the downward slope in Ca^2+^ green-5N fluorescence on the first 3 Ca^2+^ pulses. Bar graphs in panels B/D/E show means ± SE, N=5-7, with p-values (unpaired t-test) shown above error bars.

### Rewiring of glyoxal metabolism underlies cardioprotection in Alkbh7^-/-^

Several paradigms of cardioprotection against IR injury have been linked to elevated glycolysis.^27^ To test the requirement for elevated glycolysis in the protected phenotype of *Alkbh7*^-/-^, hearts were perfused in the absence of glucose (i.e., fat as the only metabolic substrate). While no difference in baseline function was observed, **Fig. S10** shows that removal of glucose abrogated cardioprotection in *Alkbh7*^-/-^. Contrary to observations with a rich substrate mix (**Fig. 4**), infarct size was significantly greater in glucose-free-perfused *Alkbh7*^-/-^ hearts vs. WT.

To further probe the requirement for elevated GLO-1 in cardioprotection, the GLO-1 inhibitor *S*-p-Bromobenzylglutathione cyclopentyl diester (SBB-GSH-CpE) was administered to hearts prior to ischemia. As **Fig. 6** shows, 1 µM SBB-GSH-CpE had no effect on WT hearts (c.f. **Fig. 4**), but completely abrogated cardioprotection in *Alkbh7*^-/-^ hearts. Separate experiments to assay GLO-1 enzyme activity in SBB-GSH-CpE treated hearts (not shown) indicated this protocol resulted in 34 ± 8% GLO-1 inhibition (mean ± SD). A significant depression of cardiac function was observed immediately upon SBB-GSH-CpE administration to *Alkbh7*^-/-^ hearts, with no effect in WT (**Fig. 6A**). Due to its higher baseline level of MGO stress (**Fig. 3D-F**), the *Alkbh7*^-/-^ heart is likely more dependent on GLO-1 activity and may therefore be hypersensitized to its inhibition. This finding suggests an important role for anti-glycation enzymes such as GLO-1 in cardiac functional homeostasis. In this regard, the mitochondrial protein DJ-1 has been shown to function as a glyoxal detoxifying enzyme,^47^ and was also recently shown to confer cardioprotection.^48,49^ However, no differences in the levels of DJ-1 were observed in *Alkbh7*^-/-^ hearts (**Fig. 6C**), suggesting that hormetic signaling in *Alkbh7*^-/-^ is somewhat specific to GLO-1 and may not engage other detoxification pathways.

**Figure 6.**
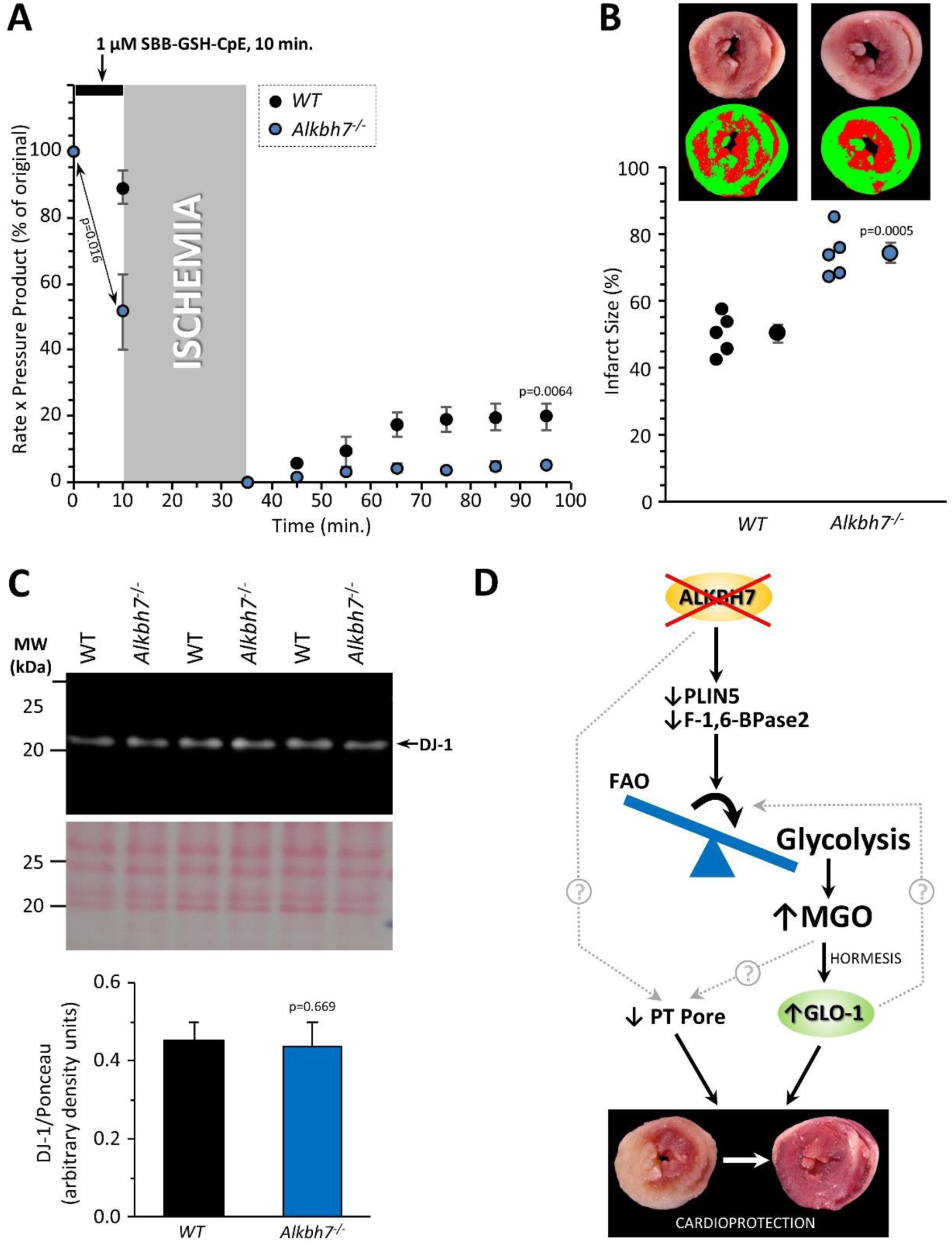
Blockade of cardioprotection in *Alkbh7*^-/-^ by GLO-1 inhibition. Hearts from young male WT and *Alkbh7*^-/-^ mice were Langendorff perfused and subjected to IR injury as in Figure 4, with delivery of 1 µM SBB-GSH-CpE for 10 min. prior to ischemia. **(A):** Cardiac function assessed by left ventricular balloon pressure transducer. Graph shows the product of heart rate multiplied by left ventricular developed pressure, as a percentage of the initial value. A significant drop in cardiac function was observed upon drug infusion in *Alkbh7*^-/-^ only (see arrow and p-value). **(B):** Post IR staining with TTC for quantitation of myocardial infarct size. Representative TTC-stained heart slices are shown, with pseudo-colored mask images used for quantitation by planimetry (red = live tissue, green = infarct). Data are quantified below, with individual data points to shown N, and means ± SE. p-values (paired t-test) are shown above error bars. **(C):** Western blot showing abundance of DJ-1 in *Alkbh7*^-/-^ and WT heart mitochondria. Ponceau stained membrane and quantitation are shown below. Bar graph shows means ± SE, N=4, with p-values (paired t-test) shown above error bars. **(D):** Schematic showing proposed events that connect loss of ALKBH7 to cardioprotection. Via mechanisms that may include downregulation of Perilipin 5 and F-1,6-BPase 2, loss of ALKBH7 causes a shift in metabolism away from fatty acid oxidation (FAO) toward elevated glycolysis. The consequent elevation in MGO leads to hormetic induction of GLO-1, which can then protect against subsequent glycative stress such as that seen in IR injury. Direct or indirect effects of ALKBH7 on the PT pore appear to play only a minor role in cardioprotection. The potential role of GLO-1 as a regulator of the metabolic shift toward glycolysis is also shown.

## DISCUSSION

Summarizing the current findings, a comprehensive analysis of the *Alkbh7*^-/-^ mouse heart suggests ALKBH7 is not a functional prolyl-hydroxylase that regulates mitochondrial activity, and that its role in necrosis involves rewiring of MGO metabolism. We propose a scheme to link these events as outlined in **Fig. 6D**. Specifically, ALKBH7 loss triggers a mild defect in fat oxidation and an elevated rate of glycolysis, possibly via perilipin 5 and fructose-1,6-bisphosphatase 2. The resulting glycative stress triggers rewiring of glyoxal metabolism, in particular GLO-1 induction, as a hormetic response which endows the added benefit of protection against IR injury. Mitochondrial PT pore opening does not appear to play a significant role in protecting *Alkbh7*^-/-^ heart. The relative position of GLO-1 as a downstream consequence of altered metabolism or as a driver of it, is highlighted by a recent systems genetic analysis which identified *Glo1* as an important gene in the regulation of lipid metabolism.^50^ While it is possible that the fat oxidation defect observed in *Alkbh7*^-/-^ mice is downstream of GLO-1, our evidence favors a hormetic response of GLO-1 to glycative stress downstream of a fat-to-glucose switch as the more likely scenario.

A surprising finding herein was that GLO-1 was upregulated in *Alkbh7*^-/-^ mice without concomitant upregulation of its companion enzyme GLO-2. The GLO-1/GLO-2 system (**Fig. 2F**) typically recycles GSH, and as predicted *Alkbh7*^-/-^ mice exhibited depleted GSH levels. While such a finding might cast focus on oxidative stress as a phenotypic driver in *Alkbh7*^-/-^, recent discoveries have implicated the GLO-1/GLO-2 system in epigenetic signaling. Specifically, the product of GLO-1, S-lactoylglutathione (SLG), has been shown to mediate the lactylation of lysine residues,^51^ including in histones,^52^ which may represent a link between metabolism and gene regulation. As such, it is possible that SLG levels may be elevated in *Alkbh7*^-/-^, which could drive epigenetic changes that underlie the phenotypes of the knockout.

The origins of the sexual dimorphism in necrosis and IR injury in *Alkbh7*^-/-^ mice remain unclear. In this regard, a recent study identified S-nitrosoglutathione reductase (GSNO-R) as a potential modulator of IR injury in male vs. female mice^53^. Notably GSNO-R can function as a formaldehyde dehydrogenase, and formaldehyde is a product of the DNA demethylation reaction carried out by many ALKBHs. As such, it is possible the role of ALKBH7 in necrosis may involve generation of formaldehyde, such that ALKBH7 deletion in males is protective by lowering the levels of this metabolite, whereas females have elevated GSNO-R levels so are already conditioned to lower formaldehyde levels.

The results herein may also provide insight to the complex biology of the diabetic heart. The incidence and progression of cardiac pathology such as heart failure is significantly worse in diabetes, and this is thought to be partly due to elevated glycative stress.^54,55^ However, somewhat paradoxically the diabetic heart is relatively protected against acute IR injury.^56-58^ As such, it is interesting to speculate whether the mechanisms of ischemic tolerance seen in the *Alkbh7*^-/-^ heart, stemming from hormetic GLO-1 upregulation, may also apply to the diabetic heart. An additional ramification of the current results may be in the area of cancer biology, where there has been interest in the potential use of GLO-1 inhibitors to target metabolic vulnerabilities of cancer cells. ^51,59^ The apparent cardiotoxic effects of SBB-GSH-CpE (**Fig. 6**) suggest that caution may be required in the use of such drugs to ensure they do not elicit cardiac toxicity.

Although the precise biologic function and substrates of the ALKBH7 enzyme remain unknown, it is tempting to speculate that the native function of the enzyme may be in the repair of glycative adducts, such that its deletion drives the responses seen herein to limit formation of such adducts. Overall, our findings highlight the importance of MGO homeostasis in the heart and suggest novel therapeutic targets for protection of tissues against IR injury. Further work is required to elucidate the signaling mechanisms that link the biochemical function of ALKBH7 to MGO metabolism.

## ACKNOWLEDGEMENTS

Work in the laboratory of PSB is funded by a grant from the US National Institutes of Health (R01-HL071158). CAK is funded by a post-doctoral fellowship from the American Heart Association (#19POST34380212). LK and EM are funded by the NHLBI-NIH Intramural Research Program (ZO1-HL002066). DF was funded by a Rochester Aging Research Center Pilot Grant. We acknowledge S. Patel and M. Gucek in the NHLBI Proteomics Core Facility for collaborating on the proteomics in murine hearts. We thank Rudi Fasan (Rochester) for help with synthesis of dimethyl-L-2HG.

## AUTHOR CONTRIBUTIONS

PSB and DF designed the studies. CAK, SMN, LK, SC and JZ performed experiments. CAK, SMN, LK, JZ, EM, DF and PSB analyzed data. HA prepared reagents. CAK and PSB wrote the manuscript with input from DF and LK, and all authors edited the manuscript and approved it for submission.

## ABBREVITATED METHODS

(for full methods see online supplement)

### Animals & materials

*Alkbh7*^-/-^ mice on a C57BL/6J background^16,17^ were bred conventionally (WT x KO), PCR genotyped at weaning, and maintained according to the “NIH Guide” (8^th^ edition, 2011) on an IACUC approved protocol. Since murine *Alkbh7*^-/-^ phenotypes are only seen in males, primarily male mice were used (except where indicated), with littermate wild-type controls, at of 8-12 weeks (young) or 1.5 years (old). All procedures were performed following administration of heparin (250 units) and tribromoethanol anesthesia (200mg/kg ip). All chemicals and reagents were from Sigma-Aldrich unless otherwise noted. Dimethyl-L-2-hydroxyglutarate was synthesized stereo-specifically^60^ and purified via silica column chromatography.

### Isolated perfused hearts

Rapidly extirpated hearts were retrograde (Langendorff) perfused at 4 ml/min. with 37°C Krebs-Henseleit buffer (KHB) gassed with 95% O_2_/5% CO_2_, and left ventricular pressure measured via a transducer-linked water-filled balloon, as described.^27^ Following 15 min. equilibration, ischemia-reperfusion (IR) injury comprised 25 min global non-flow ischemia plus 60 min. reperfusion. Hearts were then sliced and stained with triphenyltetrazoliumchloride for infarct quantitation. KHB contained 5 mM glucose, 1.2 mM lactate, 0.5 mM pyruvate and 100 µM palmitate (conjugated to BSA) as metabolic substrates, unless indicated. The following experiments were conducted: **(i)** IR alone: WT and *Alkbh7*^-/-^ hearts subjected to IR in 3 cohorts: young males, young females, old males. **(ii)** GLO-1 inhibitor IR: Young male WT and *Alkbh7*^-/-^ hearts were subjected to IR, with 1 µM SBB-GSH-CpE delivered for 10 min. prior to ischemia. A small number of SBB-GSH-CpE treated WT hearts were snap-frozen without ischemia, for measurement of GLO-1 activity. **(iii)** Glucose-free IR: Young male WT and *Alkbh7*^-/-^ hearts were perfused with KHB containing palmitate-BSA alone (no glucose, lactate, pyruvate) and subjected to IR injury. **(iv)** Proteomics & steady-state metabolomics: Young male WT and *Alkbh7*^-/-^ hearts were perfused for 15 min. then snap-frozen and stored at ^-^80°C until analysis. **(v)** Metabolic flux: Young male WT and *Alkbh7*^-/-^ hearts were perfused with KHB. Following equilibration, glucose in KHB was replaced with 5 mM [U-^13^C] glucose, and perfusion continued for 5 min., followed by snap-freezing and storage at ^-^80°C until analysis. **(vi)** Dimethyl L-2-hydroxyglutarate plus IR: Young male WT hearts were subjected to IR, with 10 µM DM-L-2-HG delivered for 20 min. prior to ischemia.

### Isolated mitochondrial experiments

Mouse heart or liver mitochondria were isolated by differential centrifugation in sucrose based media essentially as described.^61,62^ Protein was determined by the Lowry method.^63^ Mitochondrial permeability transition (PT) pore opening, induced by 100 µM CaCl_2_, was measured via spectrophotometric light scatter at 520 nm, as described.^64^ Mitochondrial Ca^2+^ handling in response to pulses of 10 µM CaCl_2_, was assayed via fluorescence of Ca^2+^-green-5N as described.^65^

### Proteomics

WT and *Alkbh7*^-/-^ hearts were perfused, snap-frozen, and shipped on dry ice. Proteomic analysis was performed essentially as described.^66^ In brief, samples were extracted in laurylmaltoside, reduced with dithiothreitol, alkylated with iodoacetamide, digested with trypsin, and peptides labeled with Tandem Mass Tag (TMT) reagents (Thermo #90110 and #A37724). Following detergent removal and desalting, samples were fractionated by basic reverse-phase HPLC (C18), and fractions were analyzed by LC-MS/MS using an Orbitrap Fusion Lumos Tribid instrument (Thermo) with an Ultimate 3000 Nano-HPLC inferface (Thermo). The instrument operated in data-dependent acquisition mode (DDA) using fourier transform (FT), scanning peptide precursors in the range 300-2000 m/z (12 ppm tolerance) and fragment ions at 100-2000 m/z (0.05 Da tolerance). Raw spectra were processed and searched within Proteome Discoverer 2.2 software (PD2.2, Thermo) using Sequest HT and Mascot algorithms and the Swiss Prot mouse database. Identified peptides were filtered for < 1% false discovery rate (FDR) using the Percolator algorithm in PD 2.2. Final lists for protein identification and quantitation were filtered by PD 2.2 with at least 2 unique peptides per protein identified with medium confidence. This method yielded 3,737 proteins with average 29% sequence coverage. Filtering for P-OH containing peptides yielded 451 peptides originating from 238 proteins. The abundance of each P-OH containing peptide was normalized to abundance of its parent protein, to quantify relative hydroxylation between WT and *Alkbh7*^-/-^.

### Identification of ALKBH7 binding partners

C-terminal FLAG-tagged human ALKBH7 was cloned into pcDNA3.1 (Invitrogen) for transient transfection (48 hr.) of HEK 293T cells. Following 1 hr. treatment with 1 mM methyl methanesulfonate (MMS) or vehicle, cell extracts were prepared as previously described.^67^ ALKBH7 interacting proteins were immunoprecipitated using anti FLAG M2 resin (Sigma), eluted with FLAG peptide, and separated by SDS-PAGE. Subsequent steps were performed by the MIT Center for Cancer Research Biopolymers Laboratory (https://ki.mit.edu/sbc/biopolymers). Excised gel bands were reduced, alkylated, trypsin-digested and desalted prior to analysis using an Agilent model 1100 Nanoflow HPLC coupled by electrospray ionization to a Thermo LTQ ion-trap mass spectrometer. Protein identification was performed using the Sequest algorithm.

### Metabolomics

WT and *Alkbh7*^-/-^ hearts were perfused, snap-frozen, pulverized, and metabolites serially extracted in 80% aqueous methanol. LC-MS/MS analysis employed a Synergi Fusion RP C18 column (Phenomenex) with an acetonitrile elution ramp, coupled to a Thermo Quantum TSQ triple-quadrupole mass spectrometer (Thermo) as previously described.^27,68^ Metabolite identification was based on retention time plus a custom SRM library built using purchased standards, with confirming fragment ions at up to 4 different collision energies (mass tolerance 0.05 Da). Data were analyzed using XCalibur Qual Browser (Thermo), with relative metabolite content normalized to the sum of all metabolites in a run.

For measurement of glycolytic flux, hearts perfused with [U-^13^C] glucose were snap frozen, extracted, and analyzed as per steady state metabolomics, with a custom SRM library used to quantify isotopologues of glycolytic metabolites. Fractional saturation with ^13^C label was calculated with correction for natural ^13^C abundance, as described previously.^27,68^

### Western blotting & blue-native electrophoresis

Hearts from male WT and *Alkbh7*^*-/-*^ mice were fractionated as previously described,^69^ and protein content determined by the Lowry method.^63^ Following SDS-PAGE and transfer to 0.2 µm nitrocellulose, membranes were probed with antibodies against ALKBH7, HSPD1, LONP1, CLPP, methylglyoxal adducts, GLO-1, HADHSC, DJ-1, and ANT1. Chemiluminescent detection (KwikQuant, Kindle Bioscience) employed HRP-linked secondary antibodies, and sample loading was normalized to Ponceau S staining of membranes immediately after transfer. Analysis of mitochondrial super-complexes by blue-native PAGE, and complex V in-gel activity assay, were accomplished essentially as described.^70^

### Cardiomyocyte isolation and Seahorse respirometry

Ca^2+^ tolerant primary adult cardiomyocytes were isolated as previously described.^61,68^ and plated at 2000/well on XF96 V3-PS plates (Agilent, Santa Clara CA). Oxygen consumption rate (OCR) was measured with a Seahorse XF96 extracellular flux analyzer (Agilent) in media containing 100 μM BSA-conjugated oleate, at baseline and following sequential injections of (i) FCCP plus oligomycin, (ii) etomoxir, (iii) antimycin A plus rotenone, for the calculation of baseline and maximally stimulated fat oxidation.

### Enzyme assays

Enzyme activities were determined in isolated mitochondria and cytosol from hearts and livers of WT and *Alkbh7*^*-/-*^ mice, as indicated. Mitochondria were freeze/thawed 3x. Complex I was measured spectrophotometrically at 340 nm as the rotenone-sensitive, coenzyme Q_1_-linked oxidation of NADH, as previously reported.^71^ Complex II was measured spectrophotometrically at 60 nm as the rate of succinate-driven, thenoyltrifluoroacetone (TTFA)-sensitive, co-enzyme Q_2_-linked reduction of dichlorophenolindophenol (DCPIP) as previously reported.^72^ α-Ketoglutarate dehydrogenase was measured spectrophotometrically at 340 nm as the α-ketoglutarate-dependent, 2-Keto-3-methyl-valerate (KMV) sensitive reduction of NAD^+^, as described.^73^ Citrate synthase was measured spectrophotometrically at 412 nm as the oxaloacetate and Acetyl CoA-linked production of 2-nitro-5-thiobenzoate (TNB) from 5,5’-dithiobis-(2-nitrobenzoic acid, DTNB).^74^ The activity of short chain and long chain specific isoforms 3-HydroxyacylCoA dehydrogenase (HADH) was measured spectrophotometrically at 340 nm as the corresponding 3-oxoacyl CoA-linked oxidation of NADH, following literature procedure.^75,76^ 3-ketopalmitoyl CoA was used for long chain HADH and acetoacetyl CoA for short chain HADH. Glyoxalase I (GLO-1) activity was measured spectrophotometrically at 240 nm as the rate of formation of *S*-D-lactoylglutathione (SLG) from the hemithioacetal adduct pre-formed *in situ* by incubation of methylglyoxal and glutathione, as reported.^77^ Glyoxalase II (GLO-2) activity was measured spectrophotometrically at 240 nm as the rate of hydrolysis of SLG, as reported.^77^

### Statistics

For all experiments, a single N (biological replicate) was considered to be the material arising from a single animal. N ranged from 3 to 17 depending on experiment, and is indicated in each figure or legend. Statistical significance was assessed by ANOVA with post-hoc Student’s t-test. Where appropriate (comparisons between littermate paired samples), paired t-tests were used. Whenever possible experiments were performed in a blinded manner.

## Footnotes

A Although ALKBH7 itself appears in the proteomic data set, this is not an indication of improper deletion. The original knockout targeted exons 2-5 containing the active site, whereas the peptides found here were in exon 1. While we cannot exclude the possibility of dominant negative effects due to an inactive truncation product, limited experiments with heterozygous animals (not shown) did not reproduce any phenotypes observed in homozygous knockouts.

