## Supplemental Information (Methods and Figures) for "ALKBH7 mediates necrosis via rewiring of glyoxal metabolism"

**Supplemental Online Information for Kulkarni *et al*. "ALKBH7 mediates necrosis via rewiring of glyoxal metabolism."**

NOTE: Original data used to generate all figures in the main document and herein are contained in a spreadsheet available at https://www.figshare.com (DOI: 10.6084/m9.figshare.12200273 reserved, unembargoed upon publication).

**CONTENTS:**

- Full Methods
- 1 Supplemental Table
- 10 Supplemental Figures
- Supplemental References

**FULL METHODS**

*Animals & materials*

*Alkbh7*$^{-/-}$ mice on a C57BL/6J background were previously described.[1,2] Mice were bred conventionally (wildtype to knockout), PCR genotyped at weaning, and were maintained according to the "NIH Guide" (8$^{th}$ edition, 2011) with food and water available *ad libitum*. Since the major phenotypes of *Alkbh7*$^{-/-}$ mice – necrotic protection and obesity – are observed in males only, male mice were used experimentally except where indicated. Littermate *Alkbh7*$^{-/-}$ and wild-type controls were used at ages of 8-12 weeks (young) or 1.5 years (old). All procedures were performed following administration of heparin (250 units) and tribromoethanol anesthesia (200mg/kg ip). All chemicals and other reagents were from Sigma-Aldrich (St. Louis MO) or VWR (Radnor PA), unless otherwise noted.

For the synthesis of dimethyl-L-2-hydroxyglutarate, a drop of concentrated HCl was added to a solution of 0.5 g (S)-(+)-5-oxo-2-tetrahydrofurancarboxylic acid (Sigma-Aldrich #301469) in 3.84 ml dry MeOH. The reaction mixture was heated to reflux and stirred vigorously overnight, then quenched with saturated $NaHCO_3$, filtered and concentrated *in vacuo*. Crude material was purified via flash chromatography on silica (50% EtOAc in n-hexane) to obtain the desired product as a clear oil, as reported.[3]

*Isolated perfused hearts*

Following heparin/anesthesia, the aorta was cannulated *in-situ* and the heart transferred to the perfusion apparatus, then retrograde perfused at 37°C with gassed (95% $O_2$/5% $CO_2$) Krebs-Henseleit buffer (KHB). Left ventricular pressure was digitally recorded at 1 kHz via a transducer-linked water-filled balloon. Following 15 min. equilibration, ischemia-reperfusion (IR) injury comprised 25 min global no-flow ischemia plus 60 min. reperfusion. After reperfusion, hearts were sliced and incubated in 1% triphenyltetrazoliumchloride for 20 min., fixed in 4% formalin for 24h., and slices digitally imaged for infarct size calculation by planimetry. In general, KHB was supplemented with 5 mM glucose, 1.2 mM lactate, 0.5 mM pyruvate and 100 µM palmitate (conjugated 6:1 with fat-free bovine serum albumin), unless modified as outlined below. The following perfusion experiments were conducted: **(i)** IR alone: WT and *Alkbh7*$^{-/-}$ hearts were subjected to IR in three separate cohorts: young males, young females, and old males. **(ii)** GLO-1 inhibitor plus IR: Young male WT and *Alkbh7*$^{-/-}$ hearts were subjected to IR as described above. After equilibration, the GLO-1 inhibitor SBB-GSH-CpE was administered at 1 µM for 10 min. prior to ischemia. No washout was used, and the drug was absent during reperfusion. A separate cohort of WT hearts were perfused with 1 µM SBB-GSH-CpE for 10 min. then freeze-clamped with Wollenberger tongs into liquid $N_2$ for later measurement of GLO-1 activity. **(iii)** Glucose-free IR: Young male WT and *Alkbh7*$^{-/-}$ hearts were perfused with KHB containing palmitate-BSA alone (i.e., no glucose, lactate or pyruvate), and subjected to IR injury. **(iv)** Proteomics & steady-state metabolomics: Young male WT and *Alkbh7*$^{-/-}$ hearts were perfused with KHB. Following 15 min. equilibration to ensure baseline contractile function, hearts were freeze-clamped with Wollenberger tongs into liquid $N_2$ and stored at $^-$80°C until analysis. **(v)** Metabolic flux: Young male WT and *Alkbh7*$^{-/-}$ hearts were perfused with normal KHB. After equilibration, glucose in KHB was replaced with 5 mM [U-$^{13}$C] glucose, and perfusion was continued for 5 min., followed by freeze-clamp with Wollenberger tongs into liquid $N_2$ and storage at $^-$80°C until analysis. **(vi)** Dimethyl L-2-hydroxyglutarate plus IR: Young male WT hearts were treated with DM-L-2-HG (10 µM final concentration) delivered from a 1000x stock solution in DMSO via a port above the perfusion cannula for 20 min., followed by IR. No washout was used, and the drug was absent during reperfusion.

*Isolated mitochondrial experiments*

Mouse heart mitochondria were isolated as previously described[4] with minor modifications. Following anesthesia, hearts were extirpated into ice-cold media comprising 300 mM sucrose, 20 mM Tris-HCl, 2 mM EGTA, pH 7.35 at 4°C. Tissue was chopped and washed twice to remove blood then homogenized in 4 ml media (IKA Tissumizer, 22,000 rpm). Homogenates were centrifuged at 800 x *g*, 5 min. Supernatants were centrifuged at 10,800 x *g*, 5 min., and pellets washed by a further 2 centrifugation steps with final resuspension in 30 µl. For $Ca^{2+}$ handling experiments (see below) the final spin utilized EGTA-free media. Protein was determined by the Lowry method.[5]

Mitochondrial permeability transition (PT) pore opening was measured as the osmotic swelling-induced decrease in light scatter at 520 nm, in a Beckman DU800 spectrophotometer. Mitochondria were incubated at 0.5 mg/ml in buffer comprising 120 mM KCl, 3 mM $KH_2PO_4$, 50 mM Tris, 5 mM succinate and 5 µM rotenone, pH 7.35 at 37˚C. Following $CaCl_2$ addition (100µM), swelling was monitored for 20 min. In some incubations, cyclosporin A (5µM) was added prior to $CaCl_2$. Mitochondrial $Ca^{2+}$ handling was assayed using the fluorescent extra-mitochondrial dye $Ca^{2+}$-green-5N. Mitochondria were incubated at 0.25 mg/ml in buffer comprising 50 mM KCl, 150 mM sucrose, 2 mM $KH_2PO_4$, 20 mM Tris, 5 mM succinate, 5 µM rotenone and 500 nM $Ca^{2+}$-green-5N, pH 7.35 at 37˚C. Pulses of 10 µM $CaCl_2$ were added every 2 min., and fluorescence measured using an Agilent Cary Varian Eclipse spectrofluorimeter ($\lambda_{EX}$ 506 nm, $\lambda_{EM}$ 530 nm).

Liver mitochondria were isolated by differential centrifugation essentially as previously described.[6] Livers were removed from anesthetized male WT and *Alkbh7*$^{-/-}$ mice and chopped into small pieces with double scissors in ice-cold liver mitochondria isolation medium (LMIM, 250 mM sucrose, 10 mM Tris hydrochloride, 1 mM EGTA, pH 7.4 at 4°C) and homogenized using a glass Dounce homogenizer. The homogenate was centrifuged at 1,000 x *g* for 3 min .and the supernatant decanted to a fresh tube, avoiding fat. This was followed by three rounds of centrifugation at 10,000 x *g*, 10 min., discarding the supernatant each time. The pellet was resuspended in 1 ml LMIM and protein quantified by the Lowry method.[5]

*Protein extraction and preparation for proteomics*

Male WT and *Alkbh7*$^{-/-}$ hearts for proteomic analysis were perfused in Langendorff mode as described above for 5 min., snap-frozen in liquid $N_2$ with Wollenberger tongs, then ground to powder and stored in two portions at -80 °C. One half of each heart was shipped on dry ice from Rochester NY to NHLBI (Bethesda MD) for proteomic analysis. Frozen heart samples were homogenized in 280 mM sucrose, 10 mM HEPES, 1 mM EGTA, and 1% (w/v) laurylmaltoside supplemented with 1X protease inhibitors (Millipore Sigma #4693159001) and 1X phosphatase inhibitors (Millipore Sigma # 4906837001) in a Precellys 24® with Cryolys (Bertin technologies). Protein was determined using a Bradford assay

(Sigma #B6916), and 100 µg protein was brought to 100 µl final volume with 100 mM triethylammonium bicarbonate (TEAB). Protein was reduced with 10 mM dithiothreitol at 55°C for 60 min. rocking at 650 rpm, then alkylated with 18 mM iodoacetamide for 60 min. protected from light. Protein was precipitated overnight with 6 volumes acetone at -20 °C, then resuspended 100 mM triethylammonium bicarbonate (TEAB) and sonicated briefly in a chilled water bath sonicator. Protein was digested with 50 µg Trypsin (Promega #V5111) overnight at 37 °C with shaking at 650 rpm. Tryptic peptides were tagged with Tandem Mass Tag (TMT) labeling reagents (Thermo Fisher #90110 and #A37724) according to the manufacturer's instructions. Labeled peptides were then lyophilized, resuspended in 50 mM ammonium bicarbonate, and layered over resin (G-Biosciences #GBS10-800) to remove residual detergent, according to manufacturer instructions. Eluted labeled tryptic peptides were lyophilized, resuspended in 0.1% (v/v) formic acid (FA), and desalted using Hydrophilic-Lipophilic-Balanced (HLB) columns (Waters #186000383) according to manufacturer instructions. Eluted peptides were lyophilized and resuspended for off-line fractionation.

*HPLC fractionation & LC-MS analysis*

Dried and labeled tryptic peptides were reconstituted with basic reverse-phase liquid chromatographic (bRPLC) buffer A (10 mM TEAB, pH 8.0) and separated using a C18 column (Xbridge 130 Å, 3.5 µm, 4.6 mm x 150mm, Waters) on a 1200 series HPLC (Agilent). The linear gradient comprised 5 – 40% solvent B (10 mM TEAB, acetonitrile, pH 8.0) over 96 minutes, with fractions collected every minute. The 96 fractions were later combined manually to 24 fractions and lyophilized.

Protein identification by LC-MS/MS employed an Orbitrap Fusion Lumos Tribid mass spectrometer (Thermo Scientific) interfaced with an Ultimate 3000 Nano-HPLC apparatus (Thermo Scientific). Peptides were fractionated by EASY-Spray PepMAP RPLC C18 column (2 µm, 100 Å, 75 µm x 50 cm) using a 120 min. linear gradient of 5 - 35% acetonitrile in 0.1% FA at 300 nl/min flow rate. The instrument was operated in data-dependent acquisition mode (DDA) using fourier transform (FT) mass analyzer for one survey MS scan. This was done on selected precursor ions followed by top 3 second data-dependent higher-energy collision (HCD)-MS/MS scans for precursor peptides with 2-7 charged ions above a threshold ion count of 10,000 with normalized collision energy of 37%. Survey scans of peptide precursors from 300 to 2000 m/z were performed at 120k resolution and MS/MS scans were acquired at 50,000 resolution with a m/z range 100-2000.

*Protein identification and analysis*

All MS and MS/MS raw spectra of TMT experiments were processed and searched using Sequest HT and Mascot algorithms within Proteome Discoverer 2.2 software (PD2.2, Thermo Scientific). Precursor mass tolerance was set at 12 ppm, fragment ion mass tolerance to 0.05 Da, trypsin enzyme with 2 mis cleavages. Carbamidomethylation of cysteine was set as a fixed modification; and TMT 6-plex (lysine), TMT 6-plex (N-term), deamidation of glutamine and asparagine, oxidation of proline and methionine were set as variable modifications. The mouse sequence database from Swiss-prot was used for database search. Identified peptides were filtered for maximum 1% false discovery rate (FDR) using the Percolator algorithm in PD 2.2 along with additional peptide confidence set to high. The final lists of protein identification and quantitation were filtered by PD 2.2 with at least 2 unique peptides per protein identified with medium confidence.

The method overall detected 49,427 peptides representing 5,642 proteins. Filtering for proteins with more than 2 peptides identified in either search engine (Mascot or Sequest) yielded 3,737 proteins with an average 29.13% sequence coverage (95% confidence interval 28.49–29.76%). Filtering the total peptide set for P-OH containing peptides yielded 625 peptides, with 451 of these having a corresponding abundance value for the parent protein, originating from a total of 238 individual proteins. The abundance of each P-OH containing peptide was normalized to abundance of its parent protein, to determine relative hydroxylation levels between WT and *Alkbh7*$^{-/-}$ paired samples.

*Immunoprecipitation to identify ALKBH7 binding partners*

The coding region for human ALKBH7 was and cloned into pcDNA3.1 (Invitrogen) for expression as C-terminal 3xFLAG tag fusion protein. Transient transfection and cellular extract production were performed as previously described.[7] Briefly, $2.5 \times 10^6$ HEK 293T cells were transiently transfected by calcium phosphate DNA precipitation with 20 μg of plasmid DNA, followed by preparation of the lysate by hypotonic freeze-thaw lysis at 48 h. post-transfection. Whole-cell extract was rotated with 10 μl of FLAG M2 antibody resin (Sigma) for 2 h. at 4 °C in lysis buffer (150 mM NaCl, 20 mM HEPES, 2 mM $MgCl_2$, 0.2 mM EGTA, 10 % (v/v) glycerol, 1 mM dithiothreitol, 0.1 mM phenylmethylsulfonyl fluoride, 0.1 % (v/v) NP-40, pH 7.9). Resin was washed extensively using the same buffer, and bound proteins eluted with two sequential volumes of wash buffer containing 100 μg/ml of 3×FLAG peptide (Sigma).

Protein identification was performed by the MIT Center for Cancer Research Biopolymers Laboratory (https://ki.mit.edu/sbc/biopolymers). Gel slices of protein bands were excised, reduced, alkylated, and digested in solution with trypsin, followed by purification and desalting of peptides on analytical C18 column tips. Peptide samples were analyzed by chromatography on an Agilent model 1100 Nanoflow high-pressure liquid chromatography (HPLC) system coupled by electrospray ionization to a Thermo LTQ ion-trap mass spectrometer. Protein identification through tandem mass spectrum correlation was performed using SEQUEST. Spectra had to match full tryptic peptides of at least 7 amino acids, have a normalized difference in cross-correlation scores (ΔCn) of at least 0.1, and have minimum cross-correlation scores (Xcorr) of 1.8 for singly charged, 2.5 for doubly charged, and 3.5 for triply charged spectra with at least 50 % ion coverage.

*Metabolomics*

Male WT and *Alkbh7*$^{-/-}$ hearts for metabolomics analysis were perfused in Langendorff mode as described above for 5 min., snap-frozen in liquid $N_2$ with Wollenberger tongs, then ground to powder and stored in two portions at -80°C. Heart powder was serially extracted in 80% aqueous methanol, extracts evaporated to dryness under $N_2$ and resuspended in 50% aqueous methanol. Liquid chromatography-tandem mass spectrometry (LC-MS/MS) analysis was performed by resolving metabolites on a Synergi Fusion RP C18 column (Phenomenex, Torrance CA) with an acetonitrile elution ramp. Metabolites were identified by retention times and by single reaction monitoring (SRM) on a Thermo Quantum TSQ triple-quadrupole mass spectrometer (Thermo Scientific, Waltham MA) as previously described.[8,9] Metabolite identification used a custom SRM library for which fragmentation patterns including confirming ions at different collision energies were empirically determined from a library of purchased chemical standards. Data were analyzed using XCalibur Qual Browser (Thermo Scientific), with relative metabolite content being normalized to the sum of all metabolites in each sample run. 8 pairs of samples were prepared and analyzed in April 2017 in a core facility setting, yielding data for 61 metabolites. A further 9 pairs of samples were prepared and analyzed in April 2019 in the senior author's laboratory, yielding additional data for 71 metabolites. Overall the analysis included 90 metabolites in total with 43 common between both data sets. As such the number of biological replicates varied between 8 and 17 depending on the metabolite in question. To process the metabolomic data set, outliers were flagged where the group-wise (WT or *Alkbh7*$^{-/-}$) data for a given metabolite exhibited a greater than 25% standard error, and individual values outside the 95% confidence intervals were removed. Missing values were imputed as weighted medians,[10] only in situations where more than 75%

of original values were still present. Of a potential 2286 total data points, 21 outliers and 32 missing values were imputed, representing 2.3% of the total data set.

For the measurement of glycolytic flux, following 20 min. of stable normoxic perfusion, $^{12}C$ glucose in KH buffer was replaced with [U-$^{13}C$] glucose, and hearts perfused for a further 5 min. Hearts were then freeze-clamped and processed similar to steady-state metabolomics, with a custom SRM library used to detect isotopomers of common metabolites. Fractional saturation of selected metabolites with $^{13}C$ label was determined, with correction for natural $^{13}C$ abundance, as described previously.[8,9]

*Western blotting*

Hearts from male WT and *Alkbh7*$^{-/-}$ mice were fractionated by differential centrifugation as previously described.[11] Protein content was determined by the Folin-Phenol (Lowry) assay.[5] Non-mitochondrial samples were diluted two-fold in SDS-PAGE sample loading buffer and incubated at 100°C for 5 min. Mitochondrial samples were diluted two-fold in sample loading buffer containing 5 times the standard concentration of SDS and incubated at 25°C for 30 min. Samples were separated by SDS-PAGE (12.5 % or 15 % gels) and transferred to 0.2 µm nitrocellulose membranes and probed with antibodies as recommended by manufacturer's protocols. Antibodies used include anti-ALKBH7 (#A2331, Abclonal, Woburn MA), anti-HSPD1 (#AP2859b Abgent, San Diego CA), anti-LONP1 (#AP19551c Abgent), anti-CLPP (#PA5-79051 Thermo-Fisher, Waltham MA), anti-methylglyoxal (#ab243074 Abcam, Cambridge MA), anti-GLO1 (#ab137098 Abcam), anti-HADHSC (#sc-376525 Santa Cruz Biotech', Dallas TX), Anti DJ-1 (#2134, Cell Signaling Technology, Danvers MA), and anti ANT1 (#ab110322, Abcam). Detection employed horseradish peroxidase-linked secondary antibodies with chemiluminescent detection (KwikQuant, Kindle Bioscience, Greenwich CT). Sample loading was normalized to Ponceau S staining of membranes immediately after transfer.

*Blue-native electrophoresis*

Mitochondrial respiratory supercomplexes were extracted and analyzed essentially as described by Beutner *et al*.[12] Briefly, mitochondria (0.5mg/ml) were incubated for 5 min. in respiration buffer comprising 120 mM KCl, 10 mM HEPES, 1 mM EGTA, 5 mM $KH_2PO_4$, 5 mM $MgCl_2$, pH 7.3 at 37˚C. After centrifugation (14000 x *g*, 10 min.) pellets were suspended in 25 µl of buffer comprising 50 mM NaCl, 40 mM imidazole, 2 mM aminocaproic acid, 1 mM EDTA, 5.7% (w/v) digitonin, pH 7 at 4˚C, and incubated on ice for 20 min. Samples were then centrifuged (14000 x *g*, 10 min.), supernatants mixed 1:1 with loading buffer (50 mM aminocaproic acid, 5% (w/v) Coomassie Blue-G), followed by resolution on 5-8% gradient blue-native gels. For the complex V in-gel assay, the gel was incubated for 2 h. in buffer comprising 35 mM Tris, 270 mM glycine, pH 8.3 at 25˚C. White precipitate complex V activity bands were visualized by adding 135 mM $MgSO_4$, 6.5 mM $Pb(NO_3)_2$, and 7.8 mM ATP to the buffer. The reaction was stopped by adding 50 % methanol and gel imaged.

*Cardiomyocyte isolation and Seahorse respirometer measurements:*

$Ca^{2+}$ tolerant primary adult cardiomyocytes were isolated from male WT and *Alkbh7*$^{-/-}$ mouse hearts by collagenase digestion as previously described,[4,9] yielding ~800,000 rod-shaped cells with >80% viability by Trypan blue assay. The final cell pellet was divided into two portions, each of which was suspended in 1 ml cardiomyocyte incubation buffer (glucose-free DMEM supplemented with 4 mM L-glutamine, 10 mM HEPES, 100 µM sodium pyruvate, 5 mM D-glucose, 500 µM L-carnitine hydrochloride, pH 7.4 at 37 $^{o}$C) and further containing either 100 µM palmitate or 100 µM oleate, conjugated to fat-free bovine serum albumin (BSA). From this suspension, cells were seeded at 2000/well on laminin-coated Seahorse XF96 V3-PS plates (Agilent, Santa Clara CA) and incubated for 1 h. in a 37 $^{o}$C humidified incubator. Cardiomyocyte incubation buffer was replaced with unbuffered DMEM (pH 7.4) containing 4 mM L-glutamine, 100 µM sodium pyruvate, 10 mM 2-deoxy-D-glucose, 500 µM L-carnitine hydrochloride and either 100 µM palmitate or 100 µM oleate conjugated to BSA. The plate was incubated for 30 min at 37 $^{o}$C, and then oxygen consumption rate (OCR) was measured with a Seahorse XF96 extracellular flux analyzer (Agilent) at baseline and following sequential injections of 1 µM FCCP + 1 µg/mL oligomycin, 5 µM etomoxir and 1 µM antimycin A + 5 µM rotenone.

*Enzyme assays*

Enzyme activities were determined in isolated mitochondria and cytosol from hearts and livers of WT and *Alkbh7*$^{-/-}$ mice as indicated. Mitochondria were freeze/thawed 3x. Complex I (NADH ubiquinone-oxidoreductase) activity was measured spectrophotometrically as rotenone-sensitive, coenzyme $Q_1$-linked oxidation of NADH as previously reported.[13] Cardiac and liver mitochondria were incubated in potassium phosphate buffer (pH 7.2) at 37 $^{o}$C containing 2.5 mg/ml BSA, 1 mM KCN, 75 µM NADH. NADH oxidation was followed at 340 nm ($\varepsilon = 6180\ M^{-1}cm^{-1}$) for 5 min. after addition of 100 µM co-enzyme $Q_1$. At the end of each run 10 µM rotenone was added and the rotenone-insensitive rate subtracted.

Complex II activity was determined spectrophotometrically as the rate of succinate-driven, thenoyltrifluoroacetone (TTFA)-sensitive, co-enzyme $Q_2$-linked reduction of dichlorophenolindophenol (DCPIP) as previously reported.[14] Cardiac and liver mitochondria were incubated in potassium phosphate buffer (pH 7.4) at 37 $^oC$ containing 120 µM DCPIP, 1 mM KCN, 10 µM rotenone, and 50 µM co-enzyme $Q_2$. The rate of DCPIP reduction was followed at 600 nM (ε = 21 $mM^{-1}cm^{-1}$) for 5 min. after addition of 5 µM succinate. At the end of each run 1 mM TTFA was added and the TTFA-insensitive rate subtracted.

α-Ketoglutarate dehydrogenase activity was determined spectrophotometrically as α-ketoglutarate-dependent, 2-Keto-3-methyl-valerate (KMV) sensitive reduction of $NAD^+$ as previously.[15] Cardiac and liver mitochondria were incubated in assay buffer comprising 35 mM $KH_2PO_4$, 5 mM $MgCl_2$, 0.5 mM EDTA, 0.05 % v/v Triton X-100, 500 µM $NAD^+$, 200 µM thiamine pyrophosphate, 40 µM reduced Coenzyme-A, 2 mM KCN, 25 µM rotenone, pH 7.25 at 37 $^oC$. The reaction was initiated by addition of 2 mM α-ketoglutarate and the rate of $NAD^+$ reduction measured at 340 nm (ε = 6220 $M^{-1}cm^{-1}$) for 5 min. At the end of each run 25 mM KMV was added and the KMV-insensitive rate subtracted.

Citrate synthase activity was determined spectrophotometrically as the oxaloacetate and Acetyl CoA-linked production of 2-nitro-5-thiobenzoate (TNB) from 5,5'-dithiobis-(2-nitrobenzoic acid, DTNB) following literature procedure.[16] Cardiac and liver mitochondria were incubated in assay buffer (100mM Tris, 0.1 % v/v Tritox X-100, 100 µM acetyl CoA, 200 µM DTNB, pH 8.0 at 37 $^oC$), the reaction initiated by addition of 200 µM oxaloacetate, and the initial linear (pre-plateau) rate of TNB formation measured at 412 nm (ε = 13.6 $mM^{-1}cm^{-1}$).

The activity of both short chain and long chain specific isoforms 3-HydroxyacylCoA dehydrogenase (HADH) was measured spectrophotometrically as the corresponding 3-oxoacyl CoA-linked oxidation of NADH, following literature procedure.[17,18] Cardiac and liver mitochondria were incubated in potassium phosphate buffer (pH 6.3) at 37 $^oC$ containing 100 µM NADH, 100 µM dithiothreitol and 0.1 % w/v Triton X-100. The reaction was initiated by addition of 50 µM 3-ketopalmitoyl CoA (for long chain HADH) or 50 µM acetoacetyl CoA (for short chain HADH) and the initial liner rate of NADH oxidation followed at 340 nm (ε = 6.22 $mM^{-1}cm^{-1}$).

Glyoxalase I (GLO-1) activity was determined spectrophotometrically as the rate of formation of *S*-D-lactoylglutathione (SLG) from the hemithioacetal adduct pre-formed *in situ* by incubation of methylglyoxal and glutathione as reported.[19] 2 mM glutathione and 2 mM methylglyoxal were incubated in 50 mM sodium phosphate buffer (pH 6.6) at 37 $^oC$ for 10 min. Cardiac or liver cytosolic extracts were then added, and the initial linear rate of SLG formation followed at 240 nm (ε = 2.86 $mM^{-1}cm^{-1}$) for 5 min. Glyoxalase II (GLO-2) activity was determined spectrophotometrically as the rate of hydrolysis of SLG, as reported.[19] 30 µM SLG was incubated in 50 mM Tris HCl buffer (pH 7.4) at 37 $^{o}$C. Cardiac or liver cytosolic extracts were added to the cuvette and the initial linear rate of SLG hydrolysis followed at 240 nm (ε = 3.10 $mM^{-1}cm^{-1}$) for 5 min.

*Statistics*

For all experiments, a single N (biological replicate) was considered to be the material arising from a single animal. N ranged from 3 to 17 depending on experiment, and is indicated in each figure or legend. Statistical significance was assessed by ANOVA with post-hoc Student's t-test. Where appropriate (comparisons between littermate paired samples), paired t-tests were used. Samples from WT and *Alkbh7*$^{-/-}$ were run in random order, and whenever possible experiments were performed in a blinded manner with the experimenter agnostic to the identity of the samples.

| *Protein* | *Accession* | *# peptides Control* | *# peptides MMS-treated* |
|---|---|---|---|
| AlkB homolog 7 | ALKB7 | 9 | 9 |
| 60 kDa heat shock protein, mitochondrial | CH60 | 3 | 10 |
| Heat shock cognate 71 kDa protein | HSP7C | 6 | 1 |
| Heat shock 70 kDa protein | HSP76 | 3 | 2 |
| T-complex protein 1 subunit ε | TCPE | 1 | 2 |
| cDNA FLJ53752, highly similar to Heat shock 70 kDa protein 1 | B4DNX1 | 2 | 0 |
| cDNA FLJ56386, highly similar to Heat shock 70 kDa protein 1L | B4DI54 | 3 | 0 |
| 10 kDa heat shock protein, mitochondrial | CH10 | 0 | 1 |
| UPF0693 protein C10orf32 | CJ032 | 0 | 1 |
| A-kinase anchor protein 12 | AKA12 | 1 | 0 |
| cDNA FLJ56389, highly similar to Elongation factor 1 γ | B4DTG2 | 1 | 0 |
| Heat shock 70 kDa protein 4 | HSP74 | 1 | 0 |
| Heat shock 70kDa protein 1A variant | Q59EJ3 | 1 | 0 |
| cDNA clone CS0DF029YN24 of Fetal brain | Q86U40 | 1 | 0 |
| RL40_HUMAN (P62987) Ubiquitin-60S ribosomal protein L40 | RL40 | 1 | 0 |
| 52 kDa Ro protein | RO52 | 1 | 0 |
| RuvB-like 1 | RUVB1 | 1 | 0 |
| Elongation factor 1-β | EF1B | 0 | 1 |
| Hemoglobin subunit β | HBB | 0 | 1 |
| NADH dehydrogenase iron-sulfur protein 7, mitochondrial | NDUS7 | 0 | 1 |
| Tubulin β-1 chain | TBB1 | 0 | 1 |
| KLRAQ motif-containing protein 1 | KLRAQ | 0 | 1 |
| cDNA FLJ36455, clone THYMU2014323, highly similar to Stabilin-1 | B3KSK0 | 0 | 1 |

**Table S1. Proteins identified as interacting with ALKBH7 by immunoprecipitation under control or MMS-treated conditions.** Cells transfected with FLAG-tagged ALKBH7 were treated with MMS as per the methods, the anti-FLAG beads used to immunoprecipitate ALKBH7-interacting proteins. Following SDS-PAGE separation, excised bands were identified by mass spectrometry of trypsin digests. Proteins are listed by name and accession, with the number of unique peptides identified under each condition indicated in the appropriate columns.

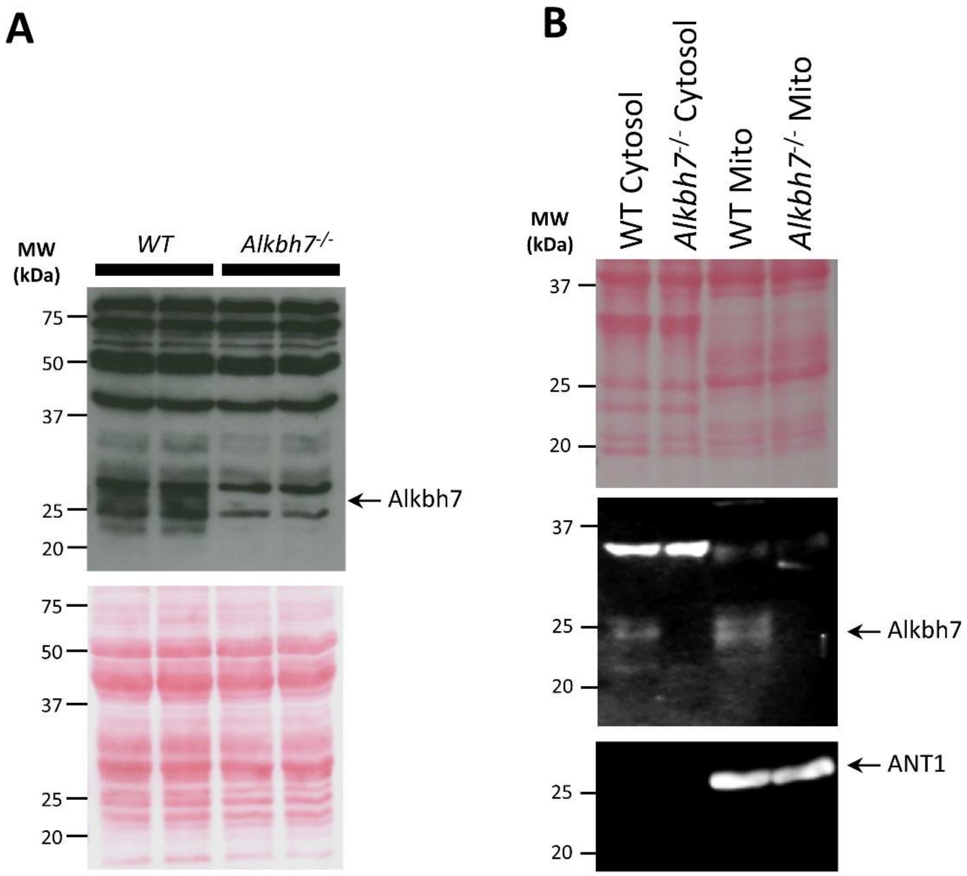


**Figure S1. Western blot showing absence of ALKBH7 protein in *Alkbh7*$^{-/-}$ mouse heart.** Hearts were fractionated into mitochondrial and cytosolic fractions by western blot. **(A):** Blot showing ALKBH7 in WT mitochondria is lost in *Alkbh7*$^{-/-}$ mitochondria. Protein loading is shown in the Ponceau S stained membrane below. Each lane represents a single animal. **(B):** Blot showing mild ALKBH7 immunoreactivity in cytosolic fraction as well as mitochondria in WT, which is lost from both fractions in the knockout. Protein loading is shown in the Ponceau S stained membrane below. Also shown is a blot for adenine nucleotide translocase 1 (ANT1), indicating no contamination of the cytosolic fraction with mitochondria. N=1 mouse per genotype. Blots are representative of at least 3 independent experiments.

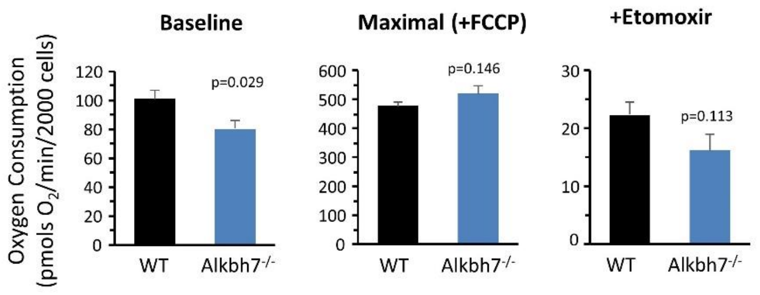


**Figure S2. Seahorse XF measurement of long chain fatty acid oxidation in WT vs. *Alkbh7*$^{-/-}$ cardiomyocytes.** Cardiomyocytes were isolated and XF measurements made as described in the methods. Graphs show oxygen consumption rate in media containing BSA-conjugated oleate as metabolic substrate. Data are shown for baseline, maximal respiration (stimulated by uncoupler FCCP in the presence of oligomycin), and in the presence of the CPT-1 inhibitor etomoxir. Note different Y-axis scales between graphs. Data are means ± SEM, N=7-9, p-values (unpaired t-test) shown above error bars.

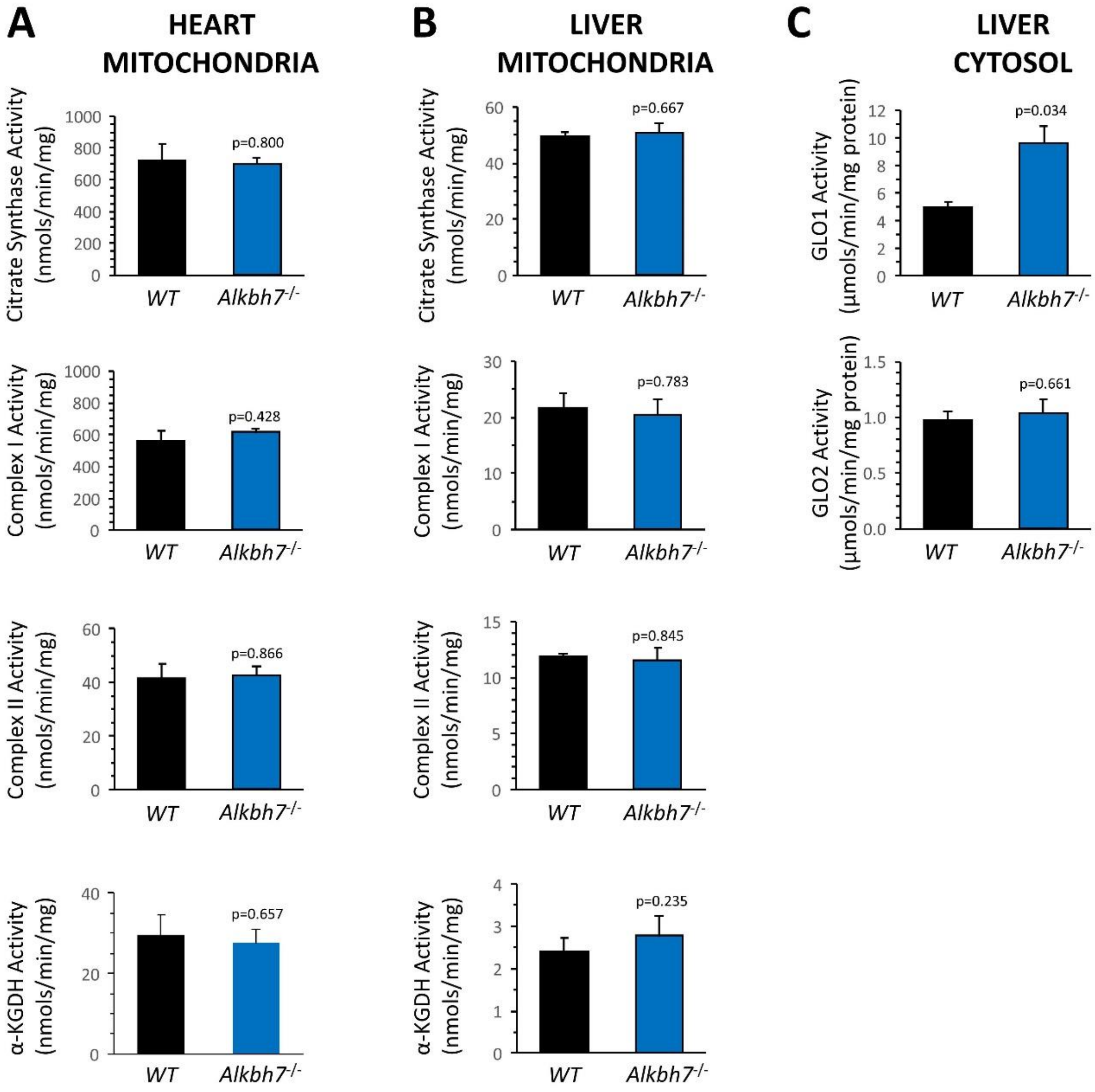


**Figure S3. Enzyme activities in WT and *Alkbh7*$^{-/-}$ tissues.** Activities of (top to bottom) citrate synthase, respiratory complex I, respiratory complex II, and α-ketoglutarate dehydrogenase, were measured spectrophotometrically as per the methods, in **(A)** Heart mitochondria, **(B)** Liver mitochondria. Panel **(C)** shows GLO-1 and GLO-2 activity in liver cytosol, complementing the heart data in Figure 2B/C. All data are means ± SEM, N=4-5, p values (paired t-tests) are shown above the error bars.

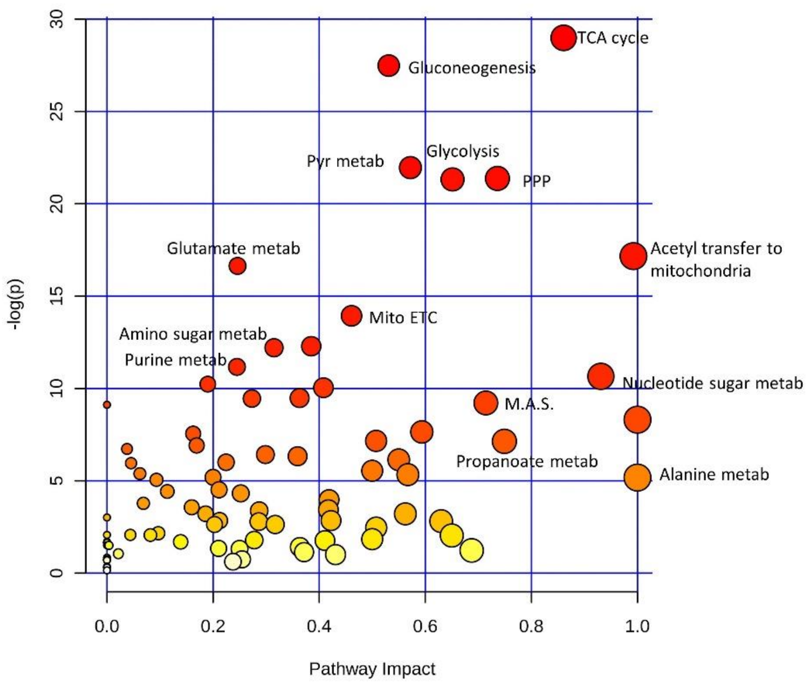


**Figure S4. Representation of metabolic pathways based on detected metabolites.** Pathway analysis was performed using open source MetaboAnalyst 3.0 software (www.metaboanalyst.ca), using all detected metabolites from the steady state analysis (Figure 3A). Greater warm coloring of a metabolic pathway indicates greater coverage, permitting conclusions to be drawn about that pathway.

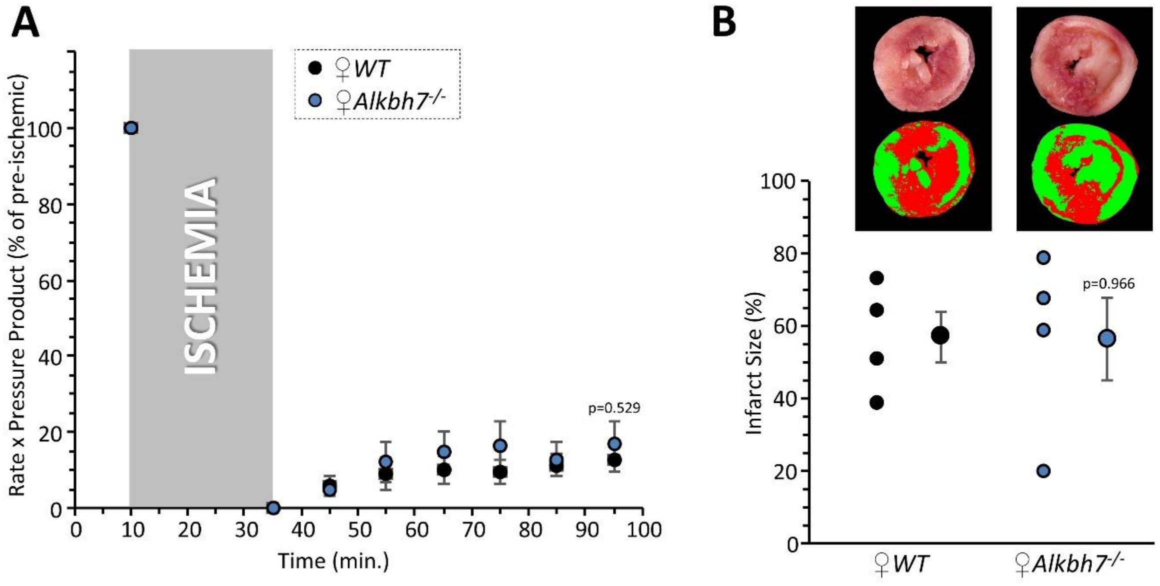


**Figure S5. Response to ex-vivo cardiac ischemia-reperfusion (IR) injury in FEMALE WT vs.** ***Alkbh7$^{-/-}$*****.** Experiments were as per Figure 4, except young female mice were used instead of male. Hearts from WT and *Alkbh7$^{-/-}$* were Langendorff perfused and subjected to 25 min. ischemia plus 60 min. reperfusion. **(A):** Cardiac function assessed by left ventricular balloon pressure transducer. Graph shows the product of heart rate multiplied by left ventricular developed pressure, as a percentage of the initial (pre-ischemic) value. **(B):** Post IR staining with TTC for quantitation of myocardial infarct size. Representative TTC-stained heart slices are shown, with pseudo-colored mask images used for quantitation by planimetry (red = live tissue, green = infarct). Data are quantified below, with individual data points to shown N, and means ± SEM. p-values (paired t-test) are shown above error bars.

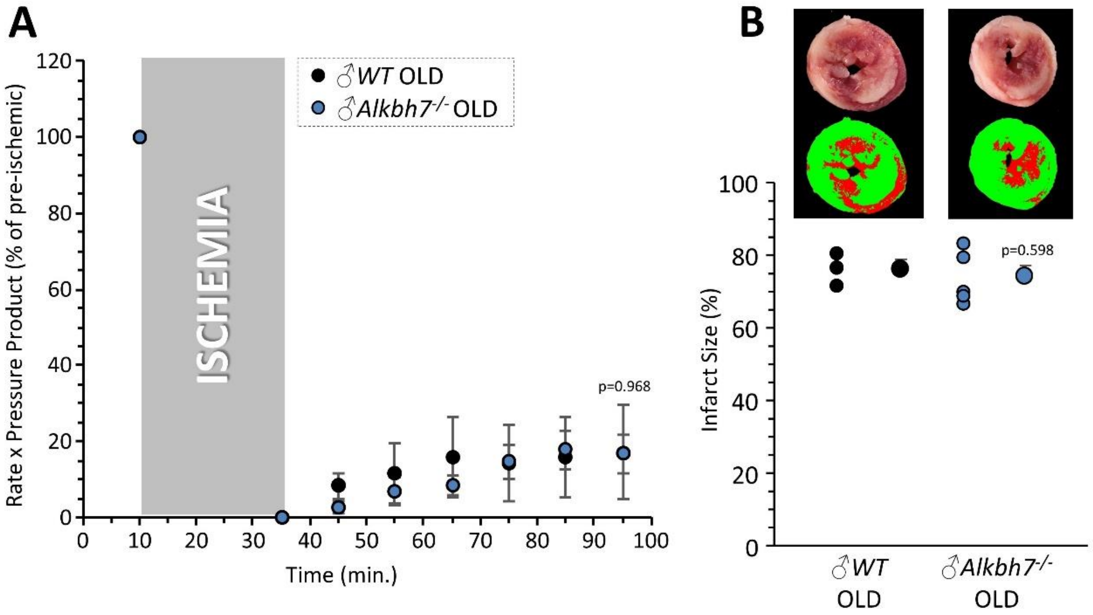


**Figure S6. Response to ex-vivo cardiac ischemia-reperfusion (IR) injury in OLD MALE WT vs. *Alkbh7*$^{-/-}$.** Experiments were as per Figure 4, except old male mice (1.5 yrs) were used instead of young males (8-12 wks). Hearts from WT and *Alkbh7*$^{-/-}$ were Langendorff perfused and subjected to 25 min. ischemia plus 60 min. reperfusion. **(A):** Cardiac function assessed by left ventricular balloon pressure transducer. Graph shows the product of heart rate multiplied by left ventricular developed pressure, as a percentage of the initial (pre-ischemic) value. **(B):** Post IR staining with TTC for quantitation of myocardial infarct size. Representative TTC-stained heart slices are shown, with pseudo-colored mask images used for quantitation by planimetry (red = live tissue, green = infarct). Data are quantified below, with individual data points to shown N, and means ± SEM. p-values (paired t-test) are shown above error bars.

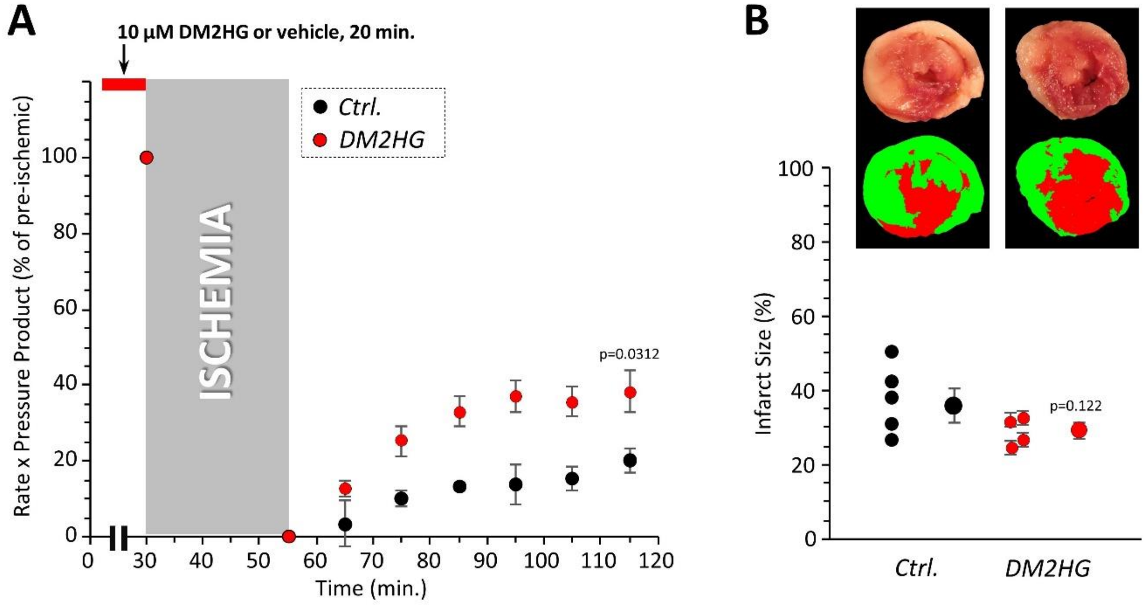


**Figure S7. Cardioprotection against IR injury by dimethyl-L-2-hydroxyglutarate.** Hearts from WT mice were Langendorff perfused and subjected to 25 min. ischemia plus 60 min. reperfusion, with optional administration of 10 µM dimethyl-L-2-hydroxyglutarate (DM2HG) for 20 min. prior to the onset of ischemia. **(A):** Cardiac function assessed by left ventricular balloon pressure transducer. Graph shows the product of heart rate multiplied by left ventricular developed pressure, as a percentage of the initial (pre-ischemic) value. **(B):** Post IR staining with TTC for quantitation of myocardial infarct size. Representative TTC-stained heart slices are shown, with pseudo-colored mask images used for quantitation by planimetry (red = live tissue, green = infarct). Data are quantified below, with individual data points to shown N, and means ± SEM. p-values (paired t-test) are shown above error bars.

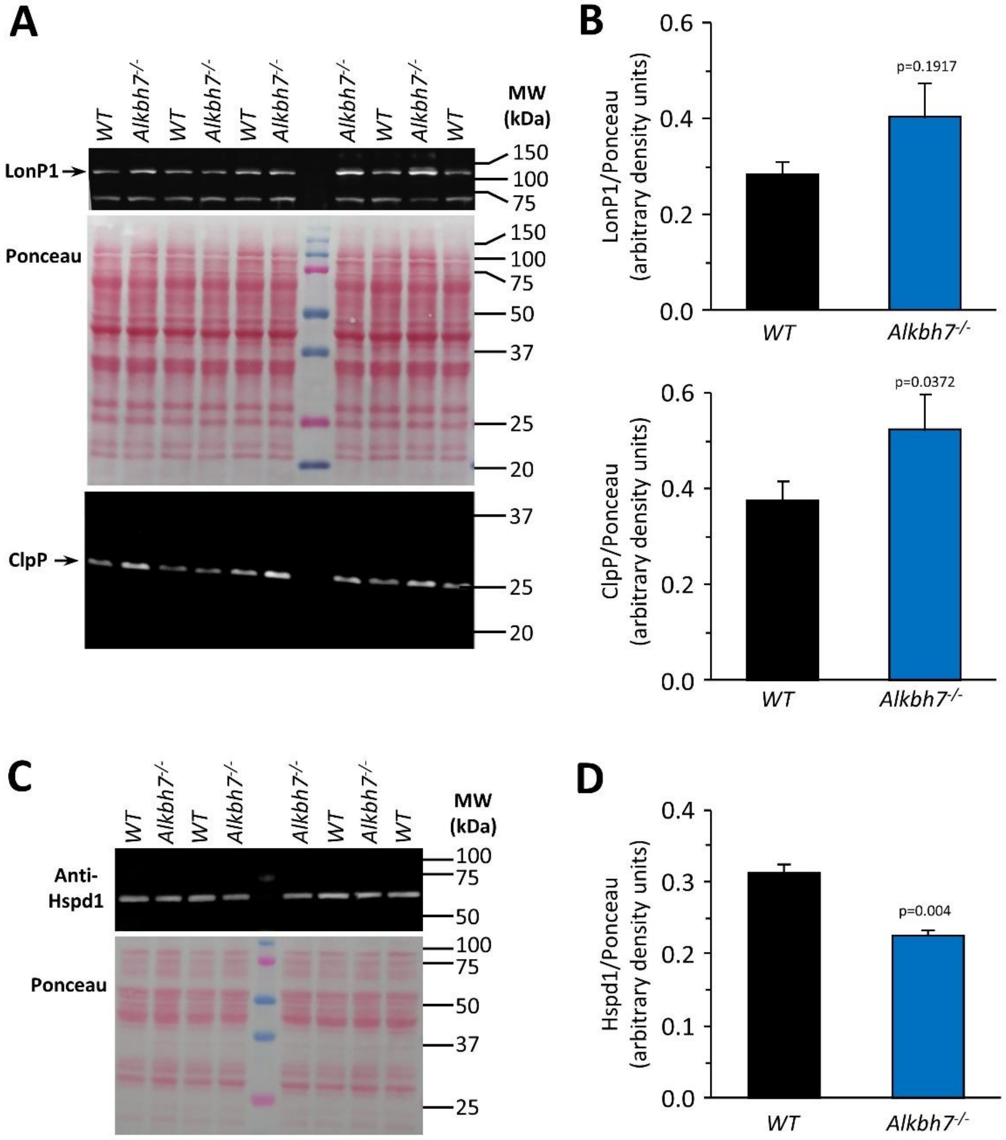


**Figure S8. Western blot detection of UPR$^{mt}$ mediators in WT vs. *Alkbh7*$^{-/-}$ hearts.** To determine -whether loss of ALKBH7 resulted in activation of the mitochondrial unfolded protein response (UPR$^{mt}$), the levels of key effectors were determined by western blot. **(A):** Blots show levels of LonP1 and Clpp in cardiac homogenate of WT and *Alkbh7*$^{-/-}$ hearts. Protein loading is shown in the central Ponceau S stained membrane. **(B):** Quantitation of data from blots of LonP1 and Clpp. Graphs show means ± SEM, N=4, with p values (unpaired t-test) above the error bars. **(C):** Blot shows level of Hspd1 (HSP60) in mitochondria from WT and *Alkbh7*$^{-/-}$ hearts. Protein loading is shown in the Ponceau S stained membrane below. **(D):** Quantitation of data from blot of Hspd1. Graphs show means ± SEM, N=4-5, with p values (unpaired t-test) above the error bars.

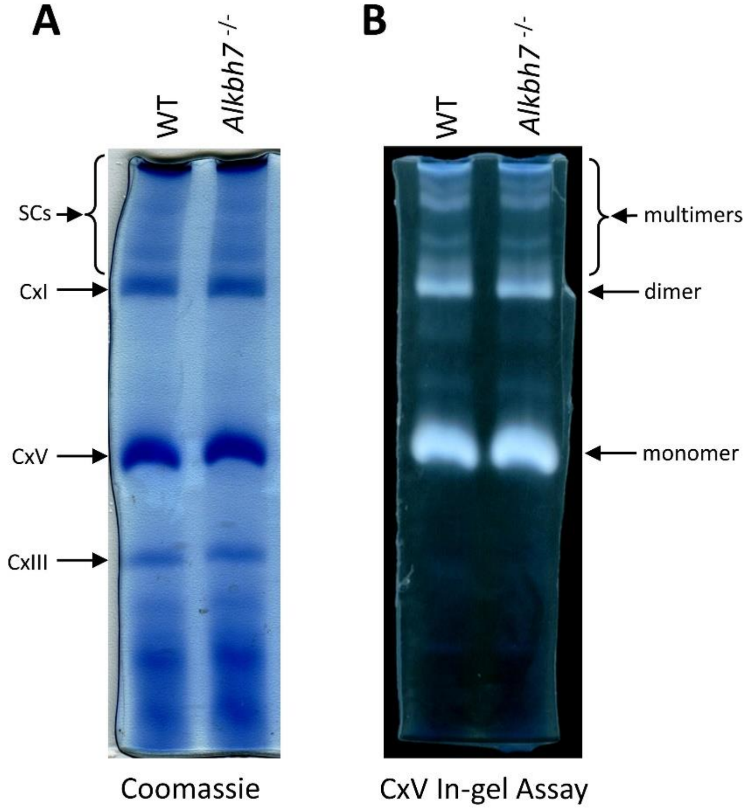


**Figure S9. Blue-native analysis of mitochondrial respiratory supercomplexes.** Heart mitochondria from WT or Alkbh7-/- mice were separated by blue-native PAGE as per the methods. **(A):** Original gel showing positions of respiratory complexes I, V and III, and supercomplexes (SCs) at the top. Note, gel was not stained post-hoc with Coomassie blue, but gel conditions contain the dye, such that gels appear with blue bands immediately upon electrophoresis. **(B):** Complex V in-gel assay. White bands show lead precipitate resulting from ATPase activity of complex V. Monomeric, dimeric, and multimeric forms of complex V are indicated. Gels are representative of at least 3 independent experiments.

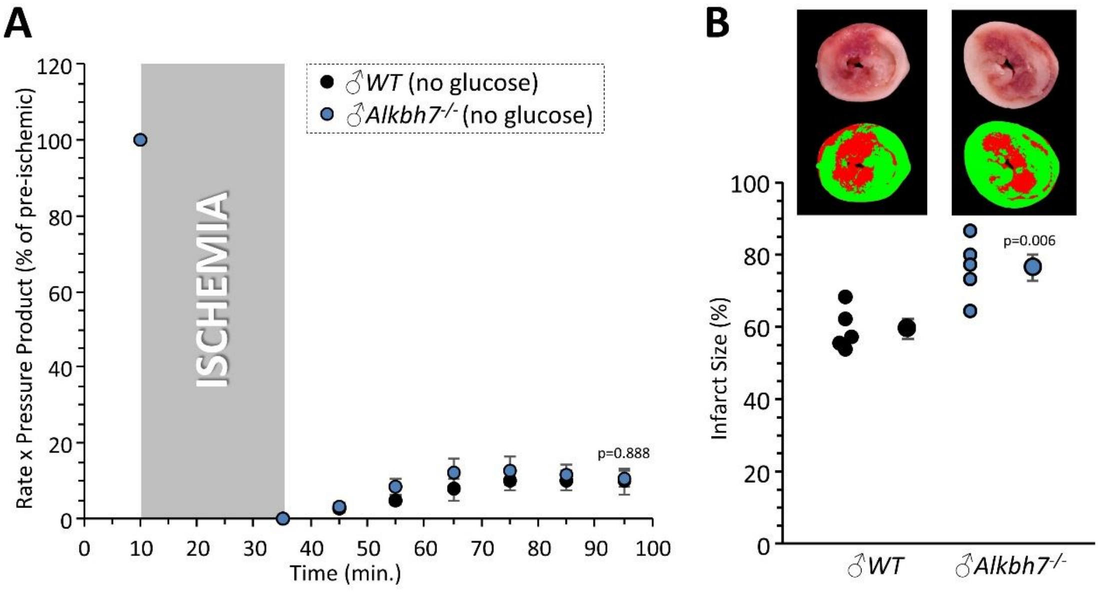


**Figure S10. Response to ex-vivo cardiac ischemia-reperfusion (IR) injury in WT vs. *Alkbh7*$^{-/-}$, in the absence of glucose.** Experiments were as per Figure 4, except that hearts were perfused with a Krebs-Henseleit buffer lacking glucose (other respiratory substrates including palmitate conjugated to BSA were still present). Hearts from WT and *Alkbh7*$^{-/-}$ were Langendorff perfused and subjected to 25 min. ischemia plus 60 min. reperfusion. **(A):** Cardiac function assessed by left ventricular balloon pressure transducer. Graph shows the product of heart rate multiplied by left ventricular developed pressure, as a percentage of the initial (pre-ischemic) value. **(B):** Post IR staining with TTC for quantitation of myocardial infarct size. Representative TTC-stained heart slices are shown, with pseudo-colored mask images used for quantitation by planimetry (red = live tissue, green = infarct). Data are quantified below, with individual data points to shown N, and means ± SEM. p-values (paired t-test) are shown above error bars.
